# An Integrated Miniaturized Metabologenomics Workflow for High-Throughput Actinomycetes Prioritization

**DOI:** 10.64898/2026.10.28.762431

**Authors:** Rima Ouchene, Laurent Falquet, Laurent Poirel, Pierre-Marie Allard

## Abstract

Antibiotic resistance is a growing global health threat, while the discovery of new antimicrobial agents remains constrained by labor-intensive cultivation, extraction, and screening workflows. We developed a miniaturized, high-throughput metabologenomics workflow integrating Duetz system-based microfermentation, extraction-free UHPLC-HRMS/MS profiling, antibacterial screening, and genome mining. A collection of 80 soil-derived actinomycetes was screened against clinically relevant pathogens, identifying seven isolates active against methicillin-resistant *Staphylococcus aureus* (MRSA), *Enterococcus faecium,* or both. All isolates were cultivated in Duetz system-based microfermentation plates, and cell-free culture filtrates were analyzed directly by UHPLC-HRMS/MS without solvent extraction. Compared with a conventional workflow, the approach reduced culture-medium consumption by approximately 99%, decreased recurring consumable costs by approximately 96%, and reduced the time required to obtain LC-MS-ready samples from typically 5 days to approximately 20 min. Integration of antibacterial, metabolomic, and genomic datasets supported the annotation of diverse metabolite families, including actinomycins, polyether ionophores, ansamycins, angucyclines, hydroxamate siderophores, ectoines, tambromycin-associated metabolites, and putative lydicamycin metabolites. Selected annotations were validated using commercial standards. Overall, this study demonstrates that integrating miniaturized Duetz system-based cultivation, extraction-free metabolomics, antibacterial screening, and genome mining provides a scalable, robust, and cost-effective platform for high-throughput strain prioritization and natural product discovery.

**Visual Abstract:** 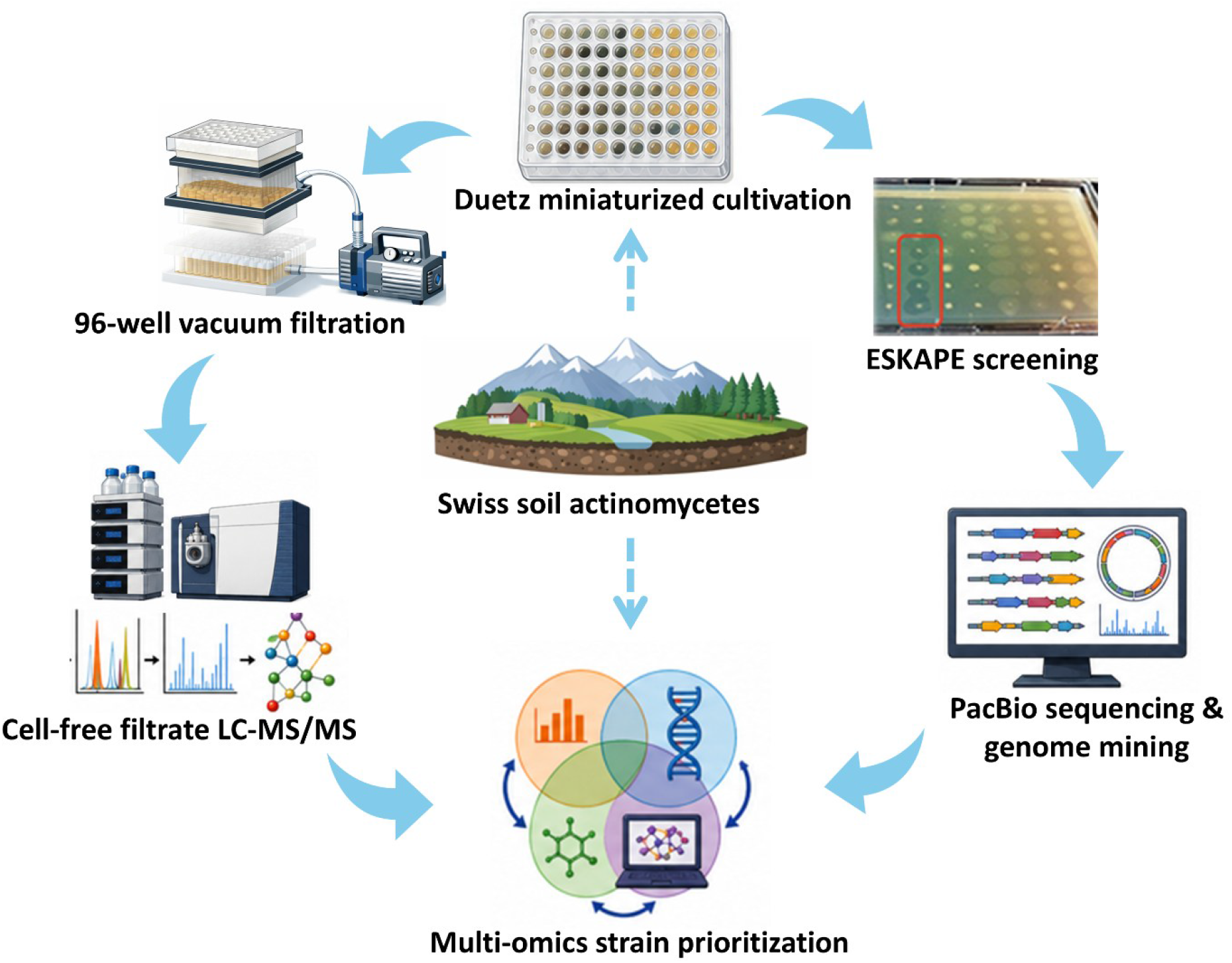

## 1. Introduction

The emergence of antibiotic-resistant microorganisms, commonly known as “superbugs”, is a major global public health concern. It has been estimated that antimicrobial resistance could cause more than 10 million deaths annually and result in cumulative economic losses exceeding USD 100 trillion by 2050.^1,2^ To guide antimicrobial research efforts, the World Health Organization (WHO) has identified several priority bacterial pathogens, including the ESKAPE pathogens: *Enterococcus faecium, Staphylococcus aureus, Klebsiella pneumoniae, Acinetobacter baumannii, Pseudomonas aeruginosa, and Enterobacter* species.^3^ Owing to their clinical relevance and elevated levels of antimicrobial resistance, five ESKAPE pathogens together with *Escherichia coli* were selected as targets for antibacterial screening in the current study.

Microbial natural products (NPs) remain a major source of antibiotics, with actinomycetes accounting for the majority of bioactive metabolites and approximately 90% of commercially available antibiotics^4^. Despite their remarkable biosynthetic potential, however, the discovery of new antibiotic scaffolds has declined substantially since the golden era of antibiotic research.^5,6^ A major challenge in contemporary natural product discovery is the efficient prioritization of strains from large and chemically diverse microbial collections.^7^ Because many strains produce previously described metabolites, screening efforts frequently lead to the rediscovery of known compounds, increasing costs and reducing discovery efficiency.^8^ Consequently, dereplication, defined as the rapid identification of known metabolites at an early stage of investigation, has become a central component of modern natural product research.^9–11^

Traditionally, strain prioritization has relied on taxonomic markers, such as 16S rRNA gene sequences, or on chemical fingerprints generated through liquid chromatography-mass spectrometry (LC-MS) profiling. Taxonomic information can support dereplication when combined with databases documenting the distribution of natural products across microbial taxa and can facilitate the selection of underexplored species for further investigation. However, substantial metabolic diversity can exist among strains sharing highly similar or even identical 16S rRNA gene sequences,^12^ limiting the predictive value of taxonomy alone. As a result, growing attention has been directed toward integrating metabolomics with functional screening approaches to obtain a more comprehensive assessment of microbial diversity and biosynthetic potential.^13–15^

Advances in sequencing technologies, bioinformatics, mass spectrometry, and metabolomics have transformed natural product discovery by enabling large-scale characterization of microbial metabolic and biosynthetic diversity.^16,8,17,18^ Metabolomics has become an indispensable tool for metabolite dereplication and strain prioritization, while genome mining provides valuable insights into biosynthetic capacity. Nevertheless, these approaches are often performed independently and typically rely on labor-intensive cultivation and solvent-based extraction procedures, making their application to large microbial collections challenging. ^19,20^ Integrating biological activity, metabolomic profiles, and genome-derived biosynthetic information within a single high-throughput workflow therefore remains a significant challenge.

To address throughput limitations, miniaturized cultivation platforms, such as the Duetz microfermentation system, have been developed for microbial growth in microtiter plates under controlled aeration.^21,22^ By enabling the parallel cultivation of large microbial strain collections using small culture volumes, the Duetz system provides an attractive platform for large-scale screening efforts. When coupled with direct UHPLC-HRMS/MS analysis of culture filtrates, it further simplifies sample preparation by eliminating conventional solvent-extraction procedures. This approach enables rapid access to metabolomic profiles from large actinomycete collections while substantially reducing sample handling, solvent consumption, and processing costs. Despite these advantages, studies combining Duetz-based microfermentation with extraction-free metabolomics remain scarce. To our knowledge, no previous investigation has integrated Duetz microfermentation, extraction-free UHPLC-HRMS/MS metabolomics, antibacterial screening against ESKAPE pathogens, and comparative genome mining within a single high-throughput strain-prioritization workflow.

In this study, we developed an integrated metabologenomics workflow combining Duetz microfermentation, antibacterial screening, direct UHPLC-HRMS/MS analysis of cell-free culture filtrates, and genome mining for the rapid exploration of actinomycete collections. Application of this approach to a collection of 80 soil-derived actinomycetes enabled biologically informed strain prioritization and supported genome-metabolome associations across the collection.

## 2. Results

### 2.1 Isolated Strains

A total of 80 actinomycete isolates were successfully recovered and selected based on colony morphotypes consistent with actinomycetes. Isolates were obtained from all sampling locations, confirming the widespread occurrence of actinomycetes across the investigated soil environments. This broad distribution ensured that the collection encompassed strains originating from diverse geographical and ecological settings. Detailed information on isolate origin, sampling sites, and geographical coordinates is provided in Supplementary Table S1 (Sheet 1).

### 2.2 Antibacterial activity of actinomycete isolates against selected ESKAPE pathogens and *Escherichia coli*

Antibacterial activity was evaluated using a spot assay against a panel of clinically relevant pathogens. Inhibition zone diameters (ID, mm) were measured as an indicator of antibacterial activity. Seven isolates (R6B, R40B, R43D, R47D, R54, R56, and R60) displayed antagonistic activity against Gram-positive pathogens. Among these, R6B, R40B, and R43D exhibited the strongest activity, producing inhibition zones exceeding 20 mm against both *S. aureus* and *E. faecium*. Isolates R54 and R56 displayed reproducible inhibition against both pathogens, with inhibition zones ranging from 15 to 21 mm, whereas R47D and R60 produced smaller inhibition zones of approximately 10-12 mm. In contrast, no inhibition was observed against *P. aeruginosa*, *E. coli*, *K. pneumoniae*, or *A. baumannii* under the conditions tested. Seven isolates displayed antibacterial activity, whereas the remaining 73 showed no measurable inhibitory effect under the conditions tested. Complete screening results are presented in Figure 1, and inhibition-zone diameters are provided in Supplementary Table S2.

**Figure 1.**
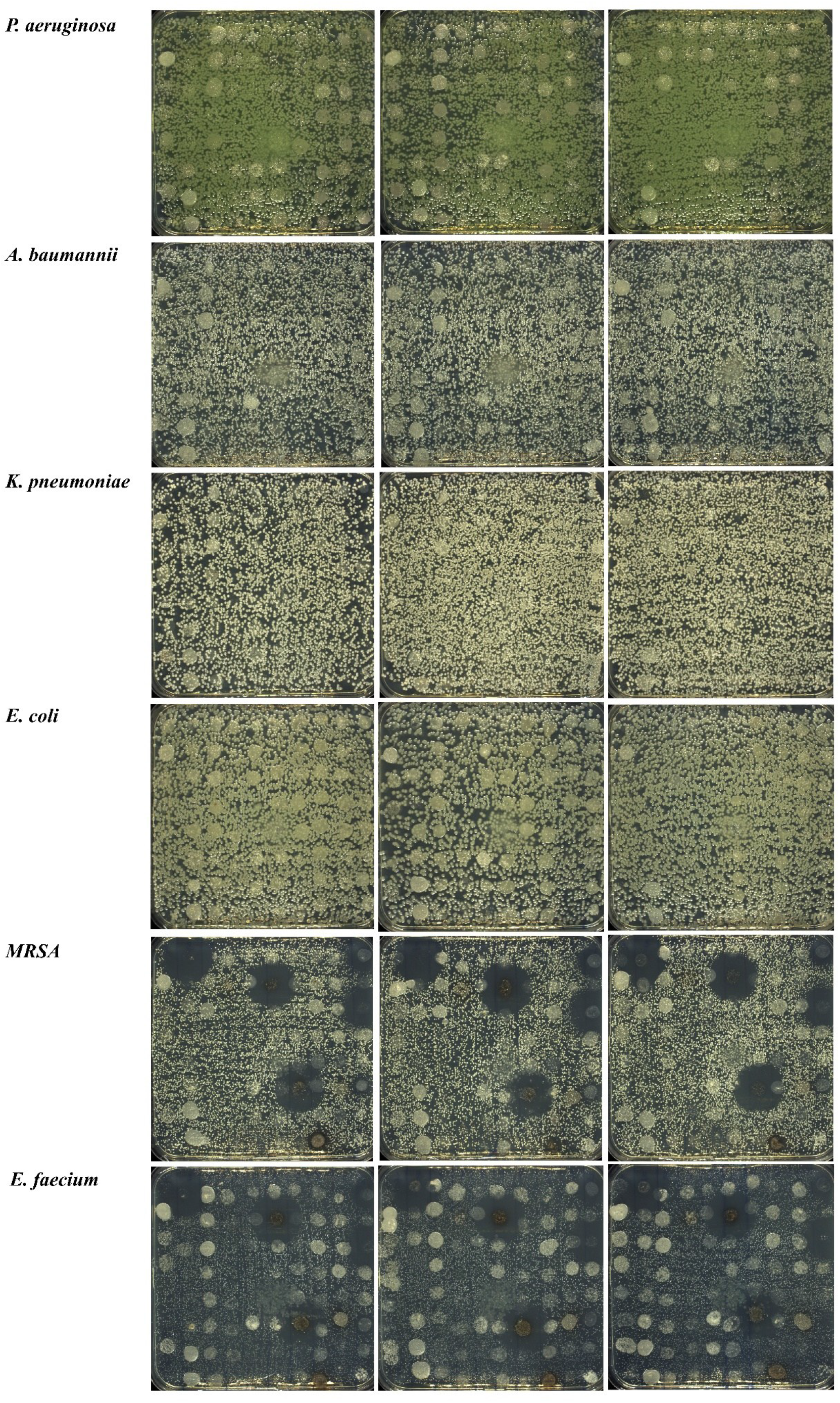
High-throughput antibacterial screening of 80 actinomycete isolates using a spot assay against clinically relevant bacterial pathogens. All assays were performed in triplicate.

### 2.3 Taxonomic assignment of sequenced isolates

Taxonomic classification revealed a clear predominance of the genus *Streptomyces* (Figure 2A; Supplementary Table S1, Sheet 2). The sequenced isolates were assigned to several *Streptomyces* lineages, including taxa affiliated with *S. albidoflavus*, *S. nigrescens*, *S. hygroscopicus*, and *S. cellulosae*. Strain R5A was the only isolate classified outside the genus *Streptomyces* and was assigned to *Spirillospora geliboluensis*.

**Figure 2.**
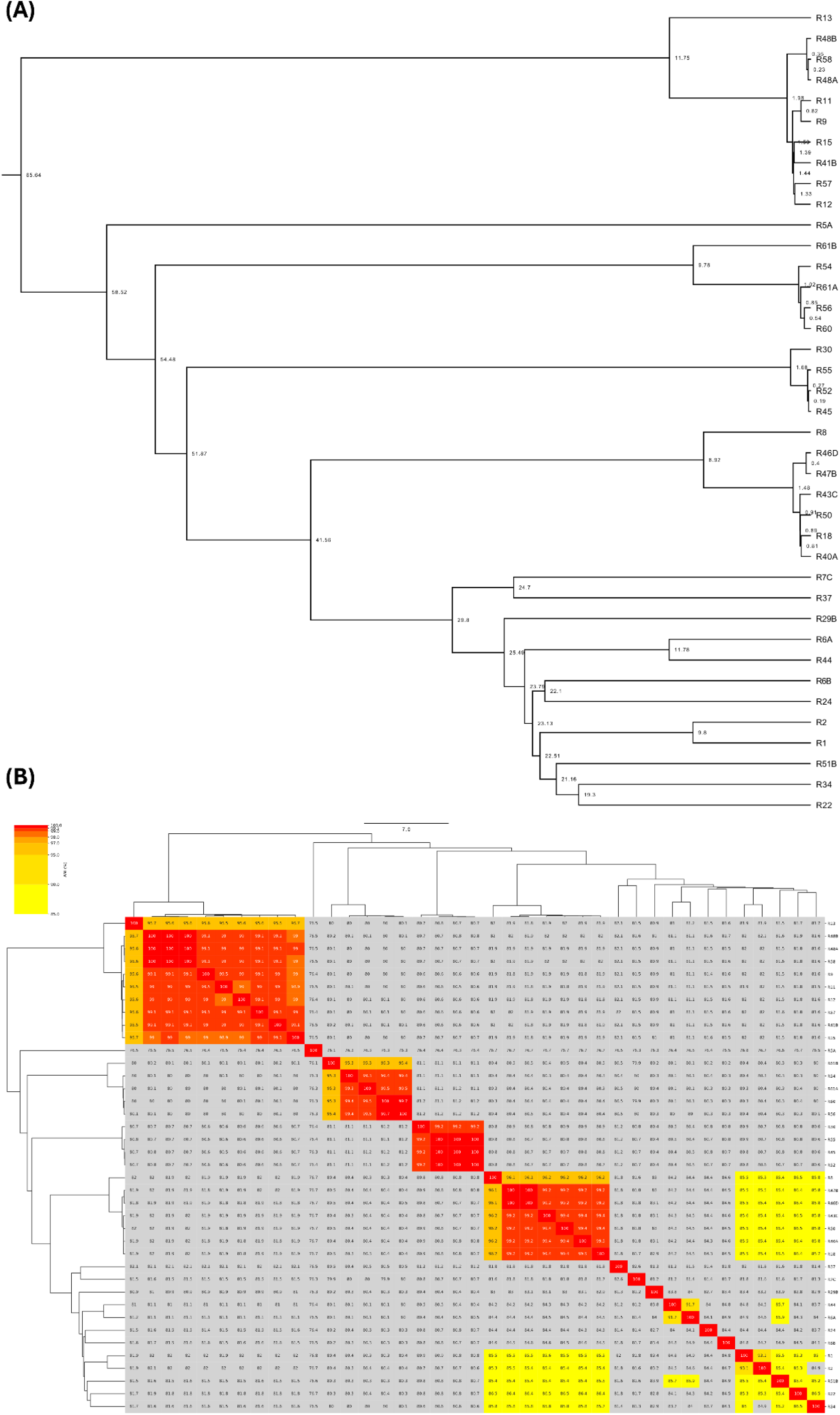
Genomic relationships among the 39 sequenced actinomycete isolates. **(A)** Maximum-likelihood phylogenomic tree inferred from the Roary core-genome alignment using IQ-TREE. Branch support values were estimated from 1,000 bootstrap replicates. **(B)** Heatmap of pairwise average nucleotide identity (ANI) values among the 39 genomes, calculated using FastANI and visualized with ANIclustermap.

Pairwise Average Nucleotide Identity (ANI) analysis revealed multiple genomic clusters within the collection (Figure 2B). Several groups of isolates displayed ANI values above 99%, indicating very closely related strains, whereas comparisons among more distant lineages generally yielded ANI values of approximately 80-85%, reflecting substantial genomic divergence. The ANI clustering was largely consistent with the phylogenomic reconstruction and highlighted the coexistence of closely related strain groups and more divergent lineages within the collection.

Overall, the collection encompassed substantial genomic diversity despite the presence of several clusters of closely related isolates. This diversity provides a relevant framework for subsequent metabologenomics analyses, enabling comparisons of metabolite production and biosynthetic potential across closely related and phylogenetically distinct strains and supporting informed strain-prioritization strategies.

The 39 genome assemblies generated in this study have been deposited in the ENA database under project accession PRJEB108104. Individual accession numbers are provided in Supplementary Table S1 (Sheet 3).

### 2.4 Genome mining and biosynthetic gene cluster prediction

AntiSMASH analysis of the 39 sequenced actinomycete genomes revealed a broad diversity of biosynthetic gene clusters (BGCs) (Supplementary Table S3). Predicted clusters included polyketide synthases (PKSs), nonribosomal peptide synthetases (NRPSs), hybrid PKS-NRPS systems, ribosomally synthesized and post-translationally modified peptides (RiPPs), terpenes, siderophores, ectoines, and other specialized metabolite classes. Several BGCs displayed high similarity to experimentally characterized pathways, whereas others showed low or no similarity to reference clusters, suggesting the presence of biosynthetic systems that remain poorly represented in currently available databases. Consistent with the genomic relationships observed in the ANI analysis, some BGC families were shared among closely related strains, whereas others appeared lineage- or strain-specific. Overall, genome mining revealed substantial biosynthetic diversity within the collection and identified numerous candidate pathways for genome-metabolome integration. Complete antiSMASH predictions for all 39 genomes are provided in Supplementary Table S3.

### 2.5 Metabolomics Analysis

Untargeted UHPLC-HRMS/MS metabolomic profiling was performed on culture filtrates from all 80 actinomycete isolates to characterize their metabolomic profiles and complement the genome mining analyses. Feature-based molecular networking (FBMN) generated a molecular network comprising 3,497 nodes, 4,761 edges, and 1,517 connected components (Figure 3). The network exhibited a clustering coefficient of 0.402 and consisted of both large interconnected molecular families and numerous isolated nodes, reflecting the substantial chemical diversity present within the collection. Molecular families varied in their distribution across the collection, with some shared among multiple strains and others restricted to individual producers. This diversity guided subsequent strain prioritization and metabolite annotation.

**Figure 3.**
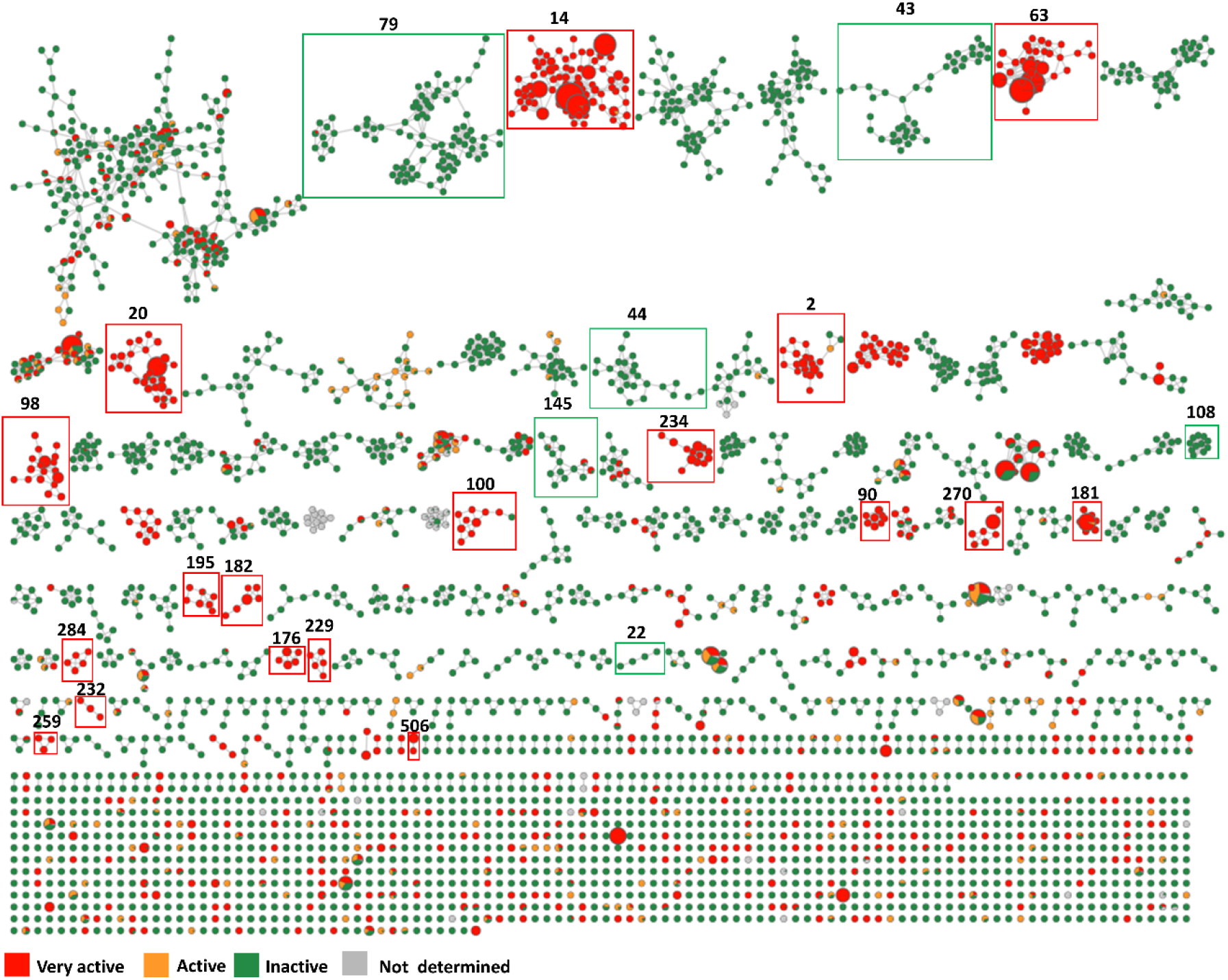
Global FBMN molecular network of the actinomycete collection. Nodes represent LC-MS/MS features, and edges represent MS/MS spectral similarity. Node size is proportional to feature abundance within the strains classified as very active. Molecular components discussed in the main text are highlighted and labeled according to their component number. Strains were classified in three groups: very active (ID ≥ 22 mm; red), active (10 mm ≤ ID < 22 mm; orange), and inactive (green).

#### 2.5.1 Multi-Omics Prioritization of Molecular Families

A comparative multi-omics prioritization strategy was applied to identify molecular families of biological, chemical, and biosynthetic interest across the actinomycete collection. Candidate molecular families were prioritized through the integration of feature-based molecular networking, GNPS spectral matching, ISDB-Lotus dereplication, SIRIUS molecular formula prediction, CANOPUS chemical classification, and antiSMASH biosynthetic gene cluster predictions. Priority was given to molecular components supported by concordant metabolomic and genomic evidence, enriched in antibacterial-active isolates, displaying strain-specific occurrence or restricted molecular subnetworks, or associated with unique and poorly characterized biosynthetic gene clusters. Similar integrated metabolome-genome approaches have recently been applied to actinomycete exploration and strain prioritization.^23^ Only molecular families fulfilling these criteria were selected for detailed investigation.

#### 2.5.2 Representative specialized metabolite families identified by FBMN and genome mining

Application of the prioritization framework described above led to the selection of several metabolite families for detailed investigation, including actinomycins, polyether ionophores, ansamycins, angucyclines, guanidyl polyene macrolides, ectoines, hydroxamate siderophores, a putative lydicamycin-like molecular family, and additional peptide-rich molecular families.

A complete list of annotated molecular features is provided in Supplementary Table S4, with features discussed in the main text highlighted in yellow. Corresponding GNPS mirror-spectrum validations are provided in Supplementary Figures S1-S7, where available. Representative chemical structures are shown, where available, to facilitate visualization of the reported metabolite families (Figures 4-13). Unless supported by additional experimental validation, structural assignments should be considered putative and based primarily on metabolomic evidence. The complete LC-MS/MS dataset, FBMN results, and associated metadata have been deposited in the MassIVE repository (<u>MSV000103110</u>).

**Figure 4.**
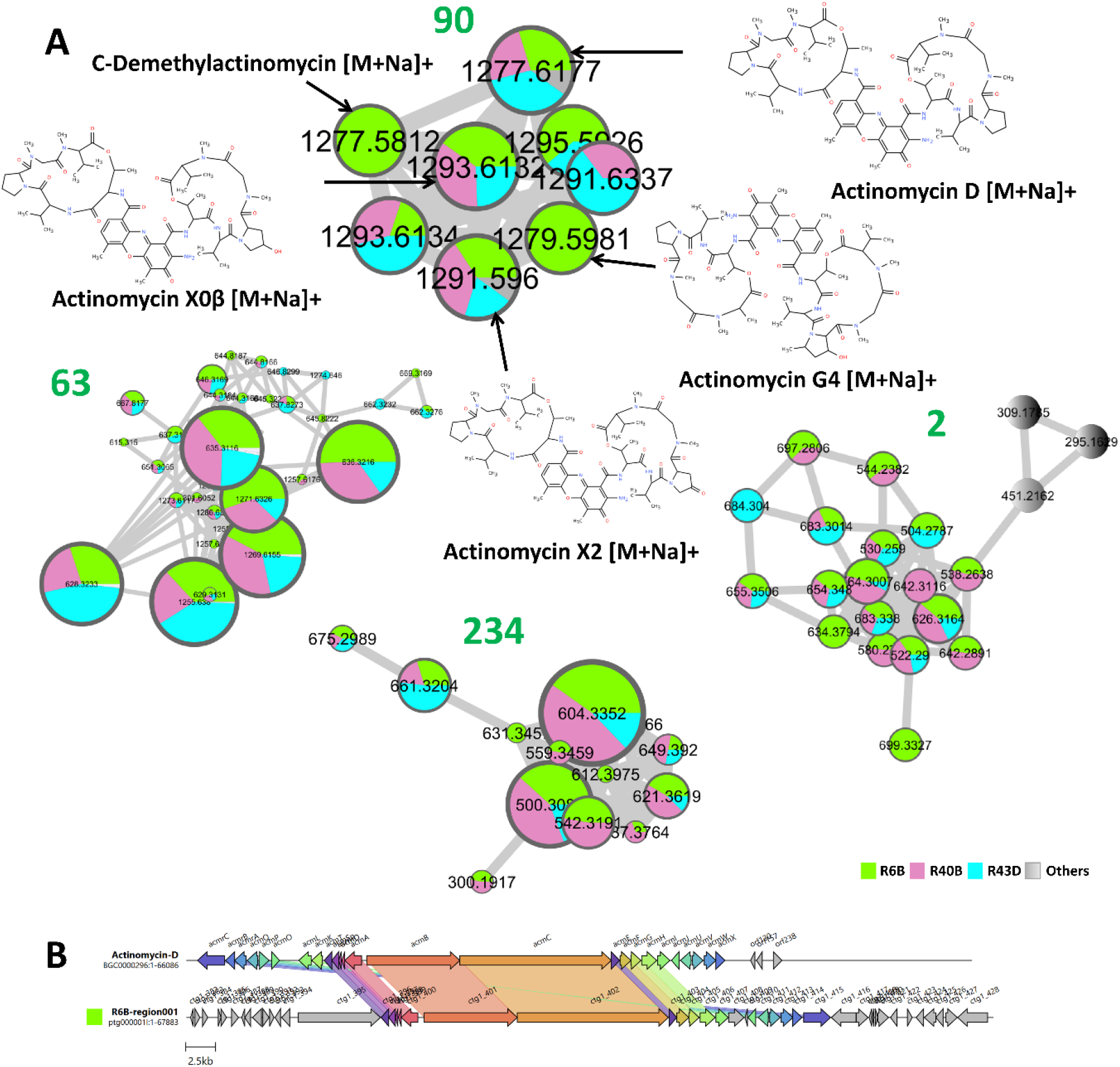
Genome-metabolome correlation of actinomycin-associated molecular families. **(A)** FBMN-derived Components 90, 63, 2, and 234 containing actinomycin-related and peptide-rich metabolites. **(B)** Comparison of the NRPS biosynthetic gene cluster identified in strain R6B with the reference actinomycin D biosynthetic gene cluster (BGC0000296).

##### Actinomycin-associated molecular families and related peptide-rich subnetworks (Components 90, 63, 2, and 234)

**Component 90** constituted the core of the actinomycin-associated molecular family. The highest-confidence annotation corresponded to <u>actinomycin D</u> (feature <u>3333</u>; *m/z* 1277.6177 [M+Na]+), which was supported by GNPS spectral library matching as well as ISDB-Lotus, SIRIUS, and CANOPUS. <u>Actinomycin X2</u> was represented by two chromatographically resolved features (<u>3325</u>; *m/z* 1291.5960 [M+Na]+ and <u>3336</u>; *m/z* 1291.6337 [M+Na]+); RT 7.44 and 7.66 min, respectively), and <u>actinomycin X0β</u>, also detected as two chromatographically resolved features (<u>3280</u> and <u>3316</u>; *m/z* 1293.6132 and 1293.6134 [M+Na]+, RT 6.74 and 7.24 min, respectively). Additional annotations included <u>actinomycin G4</u> (feature <u>3263</u>; *m/z* 1279.5981 [M+Na]+), supported by Lotus annotation and consistent classification within the actinomycin family by CANOPUS, and feature <u>3310</u> (*m/z* 1277.5812 [M+Na]+), which was consistently assigned to the actinomycin family by multiple annotation tools and tentatively annotated as a C-demethylactinomycin analogue. These metabolites were consistently classified by CANOPUS as actinomycins. Their grouping within a single FBMN component and the concordant annotations support the interpretation of Component 90 as the principal actinomycin-associated molecular family. GNPS mirror-spectrum validation further supported the annotations of actinomycin D and actinomycin X2 (Supplementary Figure S1). Comparison with a commercial actinomycin D standard provided additional support for the annotation of feature 3333 (Supplementary Table S6; Supplementary Figure S8A).

**Component 63** formed a closely connected actinomycin-enriched subnetwork, containing actinomycin D (feature <u>3330</u>; *m/z* 1255.638 [M+H]+) and actinomycin X2 (features <u>3327</u> and <u>3329</u>; *m/z* 1286.6398 [M+NH3+H]+ and 1269.6155 [M+H]+, respectively). Across the component, SIRIUS and CANOPUS consistently assigned neighboring nodes to actinomycins, oligopeptides, cyclic peptides, depsipeptides, and cyclic depsipeptides. Together, the network organization and metabolomic annotations indicate that Component 63 represents an additional actinomycin-enriched subnetwork containing known congeners and related analogues.

**Component 2** constituted an additional molecular family within the molecular network. SIRIUS and CANOPUS consistently classified most features as peptides, oligopeptides, cyclic peptides, and depsipeptides. Although features <u>1877</u> (*m/z* 504.2787) and <u>2296</u> (*m/z* 683.338) were tentatively annotated as actinomycin X0 and <u>actinoramide B</u>, respectively, the component was dominated by peptide- and depsipeptide-related metabolites, supporting its classification as a peptide-rich molecular family.

**Component 234** constituted an additional peptide-rich molecular family within the molecular network. CANOPUS consistently classified the dominant nodes as oligopeptides, peptides, depsipeptides, cyclic depsipeptides, and peptidomimetics. Representative annotations included <u>phototemtide B</u> (feature <u>2821</u>; m/z 632.3666), cyclo[Asp-Thr-Tyr-Val-Asp-Val] (feature <u>2904</u>; m/z 675.2989), and several peptide-derived metabolites. Although feature <u>2888</u> (m/z 661.3204) showed an actinomycin-like classification, the overall annotation pattern supported a peptide-rich molecular family.

Collectively, actinomycin-related metabolites were detected in strains R6B, R40B, and R43D. Comparative genome mining identified in strain R6B an NRPS biosynthetic gene cluster sharing 89% similarity with the reference actinomycin D biosynthetic gene cluster (<u>BGC0000296</u>) (Figure 4B). Combined metabolomic and genomic evidence supports the occurrence of actinomycin biosynthesis within the collection.

##### Polyether ionophore-associated molecular family (Components 14, 98, 181 and 506)

Polyether ionophore-related metabolites were detected primarily in strains R54, R56, and R60. CANOPUS consistently classified the dominant nodes from Components 14, 98, 181, and 506 within the polyether ionophore subclass. These classifications were further supported by SIRIUS molecular formula predictions and ISDB-Lotus dereplication, revealing a coherent set of annotations corresponding to known polyether ionophores and structurally related metabolites.

Within **Component 14**, several nodes were annotated as polyether ionophores. SIRIUS annotated feature <u>3341</u> (*m/z* 688.4647) as monensin, feature <u>3354</u> (*m/z* 772.522) as <u>mutalomycin</u>, and two features, including <u>3397</u> (*m/z* 742.5114, RT 8.99 min) and <u>3438</u> (*m/z* 742.5093, RT 10.00 min), both annotated as <u>nigericin</u>. Feature <u>3465</u> (*m/z* 728.4966) was annotated as <u>O-demethylabierixin</u>. The occurrence of related features with distinct retention times suggests the presence of polyether ionophore congeners or isomers. Together, the network topology and annotation patterns support the interpretation of Component 14 as a polyether ionophore-rich molecular family.

**Component 181** displayed a coherent polyether ionophore annotation pattern. Features <u>3450</u> (m/z 731.4713, RT 10.27 min) and <u>3493</u> (m/z 731.4694, RT 11.26 min) were annotated as <u>grisorixin</u> by SIRIUS. Features <u>3429</u> (*m/z* 733.4486, RT 9.75 min) and <u>3444</u> (m/z 733.4490, RT 10.18 min) received the same SIRIUS annotation corresponding to <u>O-demethylabierixin</u>, whereas feature <u>3484</u> (m/z 733.4872, RT 10.87 min) was annotated as dihydrogrisorixin. The occurrence of features with highly similar masses but distinct retention times suggests the presence of structurally related congeners or isomers. Feature <u>3478</u> (m/z 747.4637) was consistently annotated as <u>nigericin</u>. The annotation of nigericin was further supported by GNPS library matching, mirror-spectrum validation, and comparison with a commercial standard (Supplementary Table S6; Supplementary Figures S2 and S8B).

Additional support was provided by **Component 98**, in which CANOPUS consistently classified the dominant nodes as polyether ionophores. SIRIUS annotated representative features as <u>monensin</u> (feature <u>3342</u>, *m/z* 693.4188), <u>26-deoxymonensin A</u> (feature <u>3110</u>, *m/z* 677.4242), <u>mutalomycin</u> (feature <u>3355</u>, *m/z* 777.4762), and <u>O-demethylabierixin</u> (features <u>3311</u> and <u>3367</u>, *m/z* 733.4502-733.4598). Several additional nodes (features <u>3296</u>, <u>3348</u>, <u>3434</u>, and <u>3440</u>*, m/z* 719.434-719.436) displayed highly similar CANOPUS classifications and SIRIUS molecular formula predictions, suggesting the presence of closely related polyether ionophore congeners. Together, CANOPUS, SIRIUS, and ISDB-Lotus annotations support the interpretation of Component 98 as a polyether ionophore-associated molecular family. Comparison with a commercial monensin A standard further supported the occurrence of monensin-related ionophores within this component (Supplementary Table S6; Supplementary Figure S8C).

**Component 506** further supported this interpretation and consisted of two highly related features, 3401 (*m/z* 747.4638, RT 8.99 min) and 3361 (*m/z* 747.4653, RT 8.37 min), both annotated as nigericin by SIRIUS and classified as polyether ionophores by CANOPUS and ISDB-Lotus.

Several features within these molecular families were annotated as nigericin or related polyether ionophores. Comparative genome mining further revealed type I PKS biosynthetic gene clusters exhibiting 100% similarity to the reference nigericin biosynthetic gene cluster (<u>BGC0002038</u>) in strains R54, R56, and R60. Together, the recurrent detection of monensin-, mutalomycin-, grisorixin-, O-demethylabierixin-, and nigericin-like metabolites across Components 14, 98, 181, and 506 supports the presence of a coherent polyether ionophore chemical space (Figure 5B), consistent with the antibacterial activity observed against Gram-positive pathogens.

**Figure 5.**
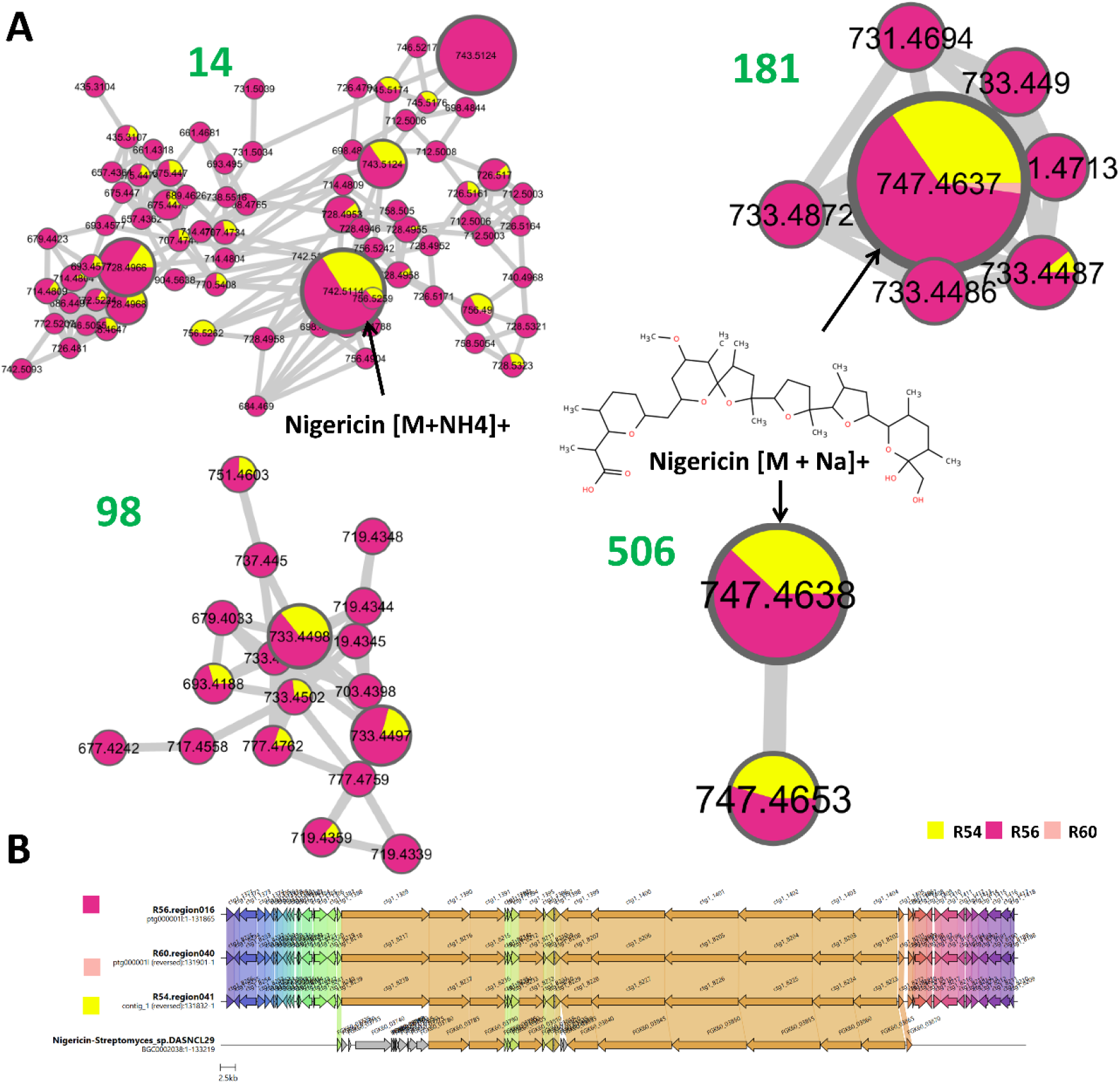
Genome-metabolome correlation of polyether ionophore-associated molecular families. **(A)** FBMN-derived Components 14, 98, 181, and 506. **(B)** Comparison of nigericin-like type I PKS biosynthetic gene clusters identified in strains R54, R56, and R60 with the reference nigericin biosynthetic gene cluster (BGC0002038).

##### Angucycline-associated molecular family (Components 108 and 22)

Angucycline-related metabolites were detected predominantly in strains R7A and R46C. FBMN analysis revealed two molecular components (108 and 22) consistently assigned to the angucycline class (Figure 6). **Component 108**, particularly abundant in strain R46C, represented the largest angucycline-rich subnetwork and was dominated by a series of closely related features centered around *m/z* 487.160. CANOPUS and ISDB consistently classified these nodes as angucyclines, while GNPS spectral matching repeatedly returned fridamycin B (e.g., features <u>2682</u> and <u>3016</u>), and SIRIUS assigned several features to <u>fridamycin A</u> (e.g., features <u>3267</u>, <u>3285</u>, and <u>3321</u>). The occurrence of multiple features sharing nearly identical masses but displaying distinct retention times suggests chromatographic isomers and/or feature over-segmentation rather than numerous distinct metabolites. Together, these annotations support the interpretation of Component 108 as a fridamycin-like angucycline molecular family. GNPS mirror-spectrum comparison further supported the annotation of feature 3016 as fridamycin B (Supplementary Figure S3).

**Figure 6.**
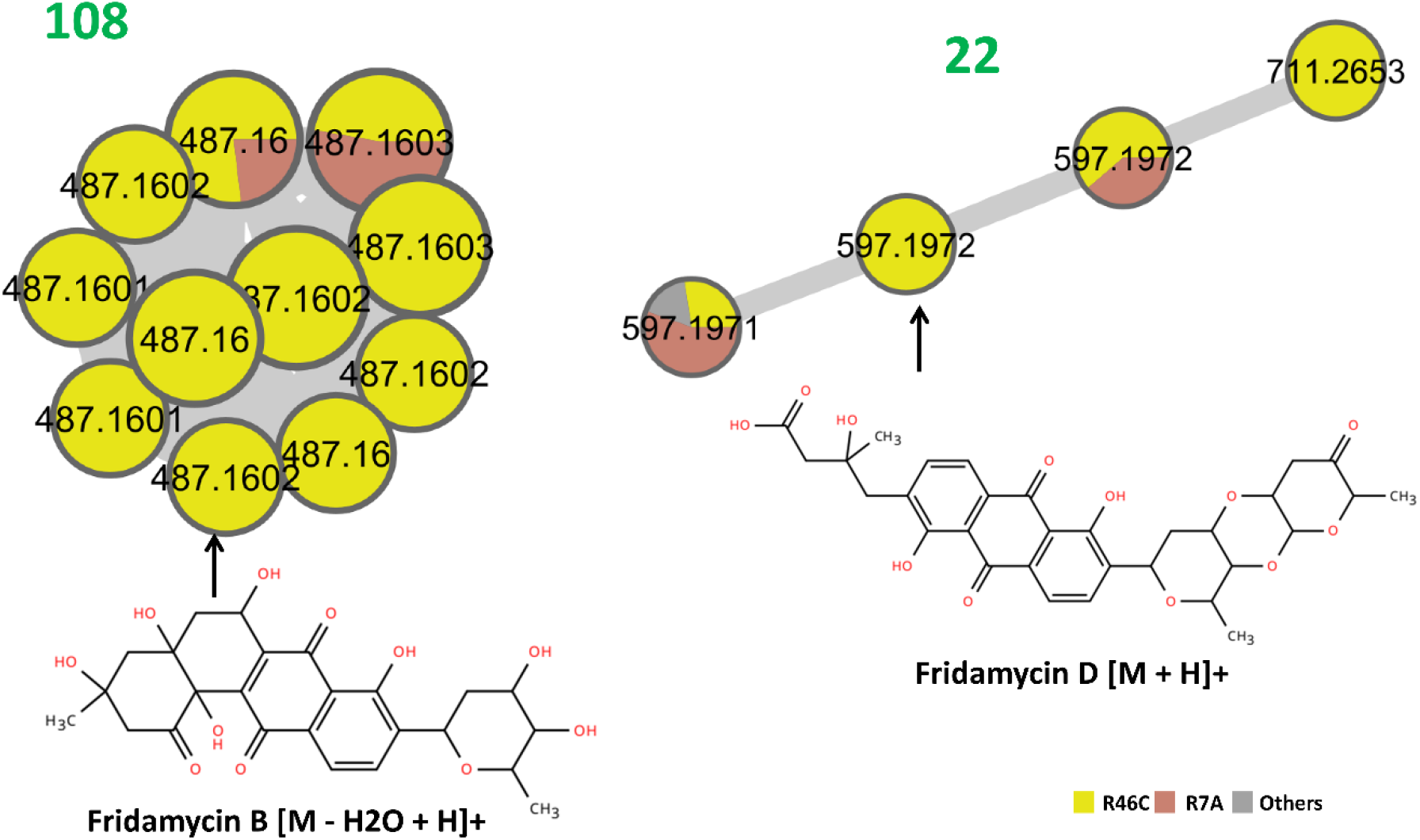
Angucycline-associated molecular families identified by FBMN (Components 108 and 22). The displayed structures correspond to fridamycin B and fridamycin D.

**Component 22** further supported this interpretation and contained several metabolites consistently classified as angucyclines by CANOPUS and ISDB-Lotus. Feature <u>3278</u> (*m/z* 597.1971) was annotated as fridamycin D by GNPS, whereas feature <u>3322</u> (*m/z* 597.1972) was annotated as fridamycin D by both GNPS and SIRIUS, whereas ISDB-Lotus suggested <u>saquayamycin B1</u>. Together, these annotations support the presence of closely related angucycline metabolites within this component.

Together, Components 108 and 22 support the presence of a diverse angucycline chemical space in strains R7A and R46C, comprising fridamycin-like metabolites centered around *m/z* 487.160 and higher-mass angucyclines represented by fridamycin D-related features 3278 (*m/z* 597.1971) and 3322 (*m/z* 597.1972).

##### Ansamycin- and ansa-macrolide-associated molecular family (Components 232, 284 and 100)

An ansamycin-associated molecular family was predominantly detected in strains R54 and R56. FBMN analysis revealed three related molecular components (232, 284, and 100) comprising metabolites consistently classified within the ansa-macrolide chemical space (Figure 7A-B).

**Figure 7.**
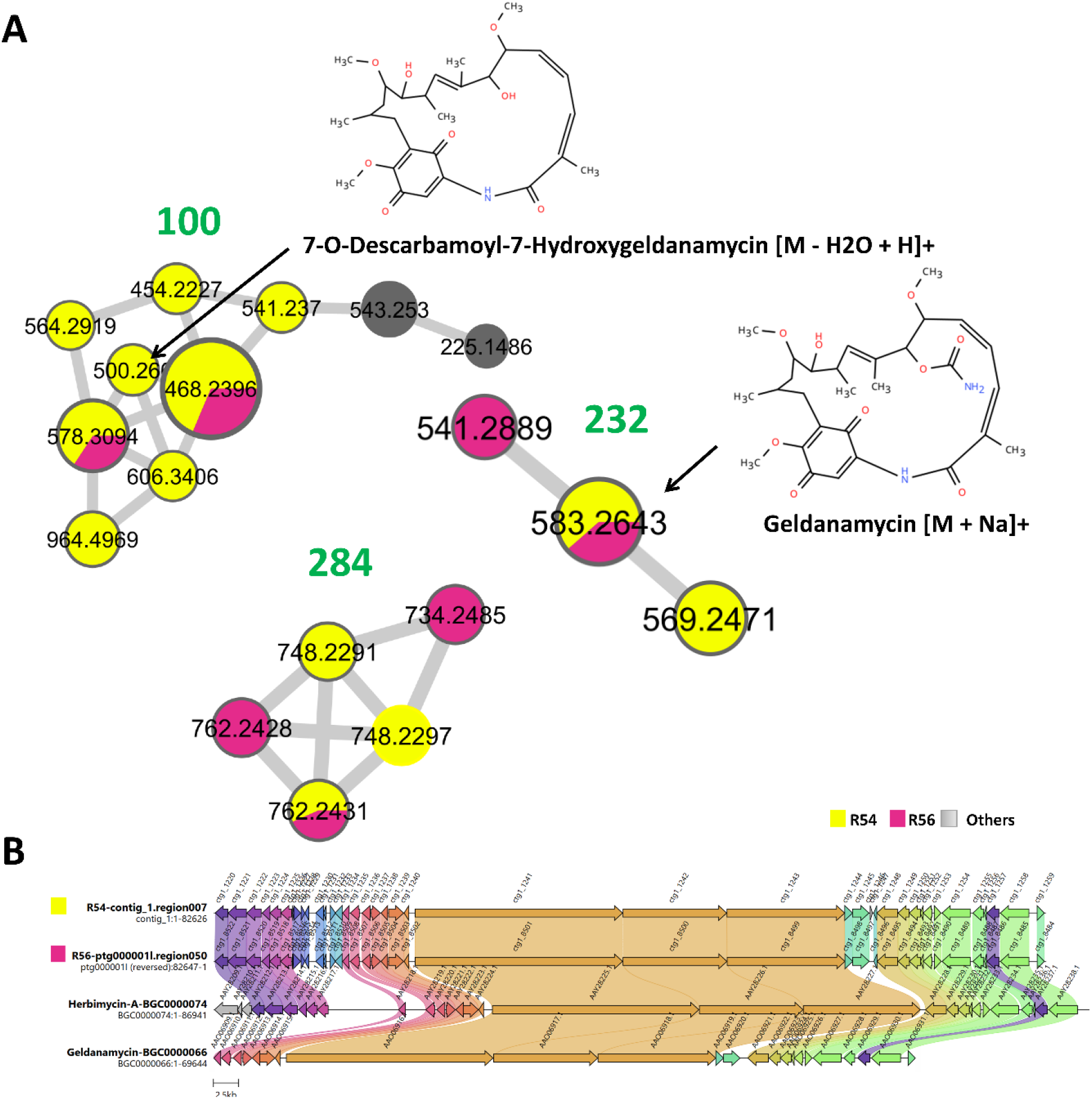
Genome-metabolome correlation of the ansamycin-associated molecular family. **(A)** FBMN-derived Components 232, 284, and 100. **(B)** Comparison of biosynthetic gene clusters identified in strains R54 and R56 with the reference herbimycin A (BGC0000074) and geldanamycin (BGC0000066) biosynthetic pathways.

**Figure 8.**
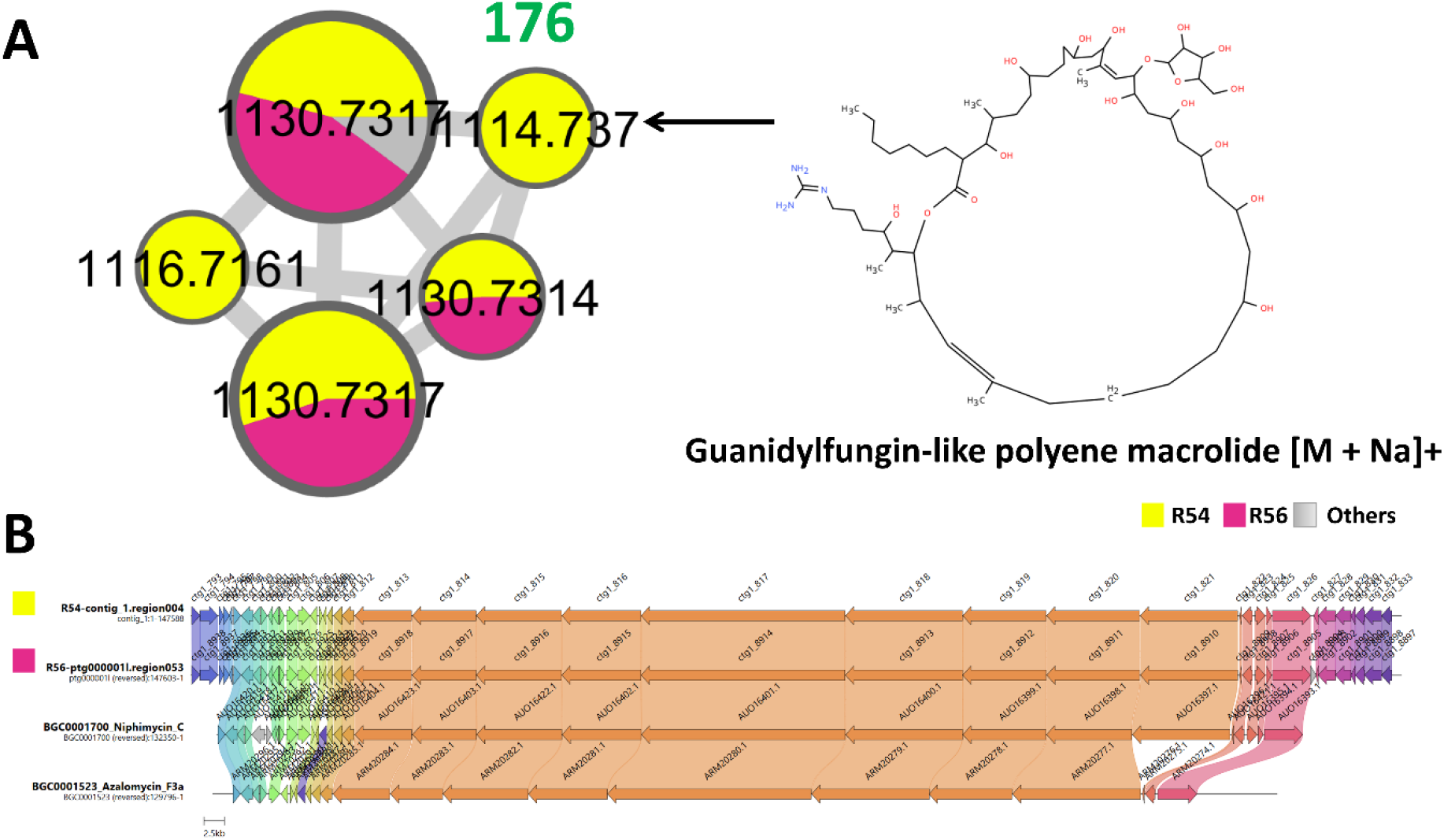
Genome-metabolome correlation of a guanidylfungin-like polyene macrolide molecular family. **(A)** FBMN-derived Component 176. **(B)** Comparison of type I PKS biosynthetic gene clusters identified in strains R54 and R56 with the reference Niphimycin C-E (BGC0001700) and Azalomycin F3a (BGC0001523) biosynthetic pathways.

**Component 232** provided strong evidence for the presence of <u>ansamycin</u>-type polyketides. Feature <u>3230</u> (*m/z* 583.2643) was annotated as <u>geldanamycin</u> by both GNPS and SIRIUS (cosine score = 0.915; Supplementary Figure S4). Additional connected nodes included feature <u>2445</u> (*m/z* 541.2889), dereplicated as a <u>macbecin</u>-related metabolite, and feature <u>3028</u> (*m/z* 569.2471), assigned to a structurally related ansamycin-like derivative. All three features were consistently classified as ansa-macrolides by CANOPUS and ISDB-Lotus. Together, these annotations support the interpretation of Component 232 as an ansamycin-rich molecular family.

Additional support for the ansa-macrolide classification was provided by Component 284. While feature <u>1668</u> (*m/z* 734.2485) showed a tentative ISDB-Lotus match to Rifamycin S and CANOPUS classified the component as an ansa-macrolide, no confident annotations were obtained for the remaining nodes.

Additional support for the ansa-macrolide chemical space was provided by Component 100. Feature <u>3234</u> (*m/z* 500.2660) was annotated as <u>7-O-descarbamoyl-7-hydroxygeldanamycin</u>, while several additional nodes were classified as macrolides or ansa-macrolides. Together, these annotations support the association of Component 100 with the broader ansamycin-related chemical space.

The recurrence of geldanamycin-, macbecin-, and related ansa-macrolide annotations across Components 232, 284, and 100, together with consistent CANOPUS assignments to macrolide and ansa-macrolide chemical classes, supports the presence of an ansamycin-associated chemical space in strains R54 and R56. Comparative genome mining further reinforced this interpretation, revealing in strain R54 a type I PKS biosynthetic gene cluster displaying 90% similarity to the herbimycin A biosynthetic gene cluster (<u>BGC0000074</u>) and substantial similarity to the closely related geldanamycin biosynthetic pathway (<u>BGC0000066</u>) (Figure 7B). Taken together, the metabolomic and genomic data support the occurrence of geldanamycin-and macbecin-related ansamycins in these strains, whereas the rifamycin-related annotation remains tentative and is supported only by metabolomic evidence.

##### Guanidylfungin-like polyene macrolide molecular family (Component 176)

Feature <u>2948</u> (*m/z* 1114.737) was classified as a polyene macrolide polyketide by SIRIUS and CANOPUS. Associated features <u>2908</u> (*m/z* 1116.7161), <u>2921</u>, <u>3022</u>, and <u>3053</u> (*m/z* 1130.731) formed a closely related cluster and yielded recurrent guanidylfungin-like annotations, including <u>guanidylfungin B</u> for feature 2908. Together, these annotations support the occurrence of a guanidylfungin-related polyene macrolide molecular family.

Comparative genome mining further supported this interpretation. Highly similar Type I PKS biosynthetic gene clusters were identified in strains R54 and R56, both displaying 87% similarity to the <u>niphimycin C</u> biosynthetic pathway (<u>BGC0001700</u>) and substantial synteny with the related <u>azalomycin F3a</u> pathway (<u>BGC0001523</u>). Given the close biosynthetic relationship among niphimycins, azalomycins, and guanidylfungins, the concordance between metabolomic and genomic data supports the occurrence of guanidylfungin-like polyene macrolides in strains R54 and R56.

##### Ectoine-associated molecular family (Component 145)

Component 145 was predominantly detected in inactive strains (Figure 9A-B). Features <u>15</u> (*m/z* 159.0766), <u>83</u> (*m/z* 159.0764), and <u>555</u> (*m/z* 159.0765), which were consistently annotated as <u>5-hydroxyectoine</u> by GNPS. The occurrence of multiple closely related features assigned to 5-hydroxyectoine likely reflects feature splitting during peak detection and alignment rather than distinct metabolites. ISDB-Lotus and CANOPUS further classified these features within amino acid-related chemical classes, providing additional support for their assignment. Additional support was provided by feature <u>214</u> (*m/z* 143.0816), annotated as ectoine by GNPS library matching. GNPS mirror-spectrum comparisons (Supplementary Figure S5) and analyses of commercial ectoine and 5-hydroxyectoine standards (Supplementary Table S6; Supplementary Figures S8E-F) further confirmed these annotations.

**Figure 9.**
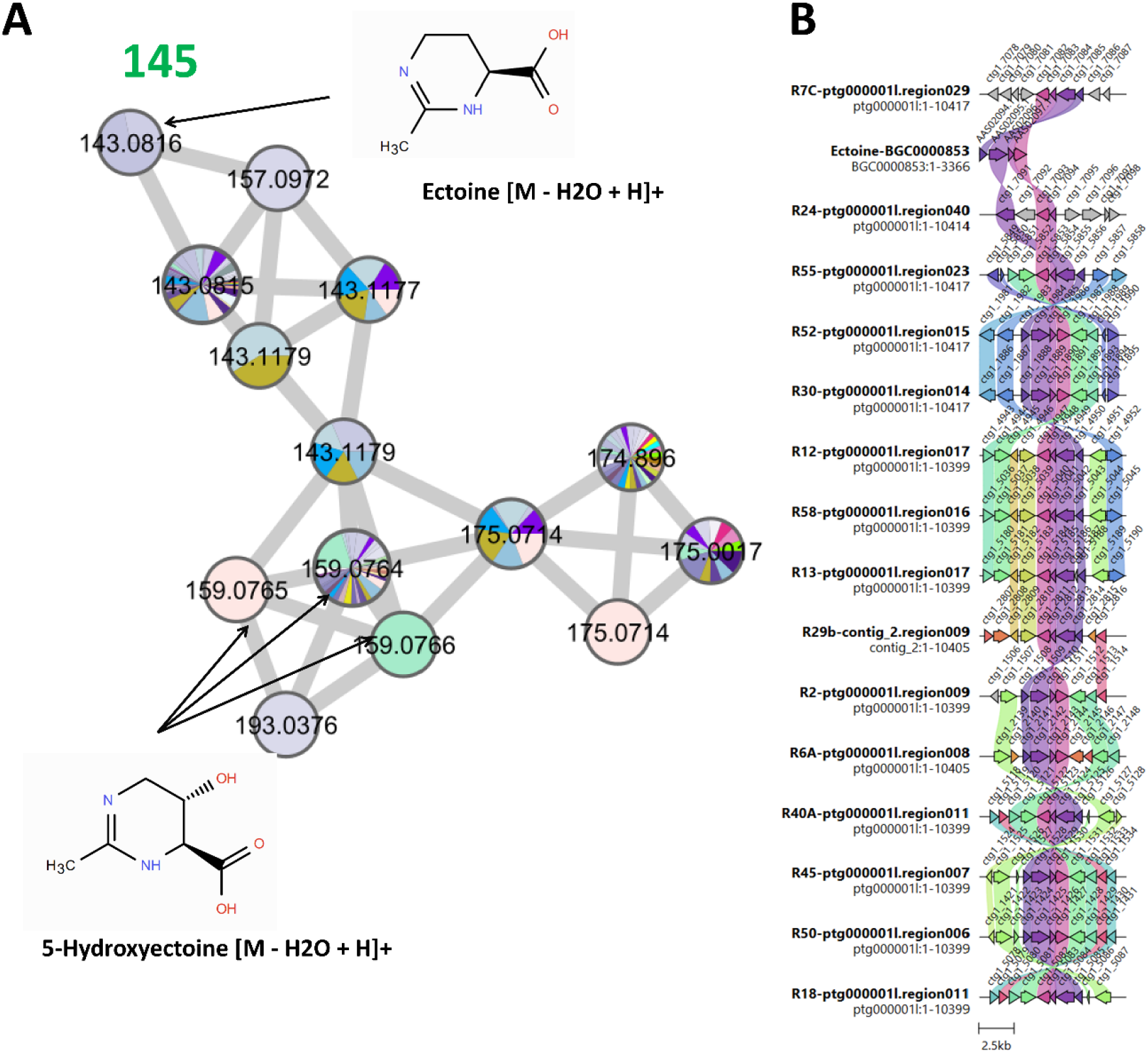
Genome-metabolome correlation of the ectoine-associated molecular family. **(A)** FBMN-derived Component 145. The displayed structures correspond to ectoine and 5-hydroxyectoine. **(B)** Ectoine biosynthetic gene clusters identified in the strain collection compared with the reference ectoine biosynthetic pathway (BGC0000853).

Comparative genome mining further supported these findings. Highly conserved ectoine biosynthetic gene clusters were identified in 36 of the 39 sequenced isolates (Figure 9B), displaying strong conservation of gene content and synteny relative to the reference ectoine biosynthetic pathway (<u>BGC0000853</u>). Together, the metabolomic and genomic evidence supports the widespread production of ectoine and 5-hydroxyectoine across the strain collection.

##### Lydicamycin-related molecular family (Component 79)

A putative lydicamycin-like molecular family (**Component 79**) was detected predominantly in strains R29A, R30, R31, R45, R51A, R52, and R55 (Figure 10A-B). The high-mass region of the network was centered on feature <u>2501</u> (*m/z* 841.5328), which was annotated by GNPS as the <u>lydicamycin</u> derivative TPU-0037A. Additional highly connected features, including <u>2482</u> and <u>2574</u> (*m/z* 783.4908-783.4898), showed tentative similarity to bafilomycin-related macrolides based on ISDB-Lotus; however, these annotations were not consistently supported across the molecular family. In contrast, neighboring nodes <u>2618</u> (m/z 797.5068), <u>2633</u> (*m/z* 797.5065), <u>2612</u> (*m/z* 811.5222), and <u>2777</u> (*m/z* 825.5379) formed a coherent series of closely related high-mass congeners. Together, the network organization and GNPS annotations support the occurrence of a lydicamycin-related molecular family, further supported by mirror-spectrum comparison (Supplementary Figure S6).

**Figure 10.**
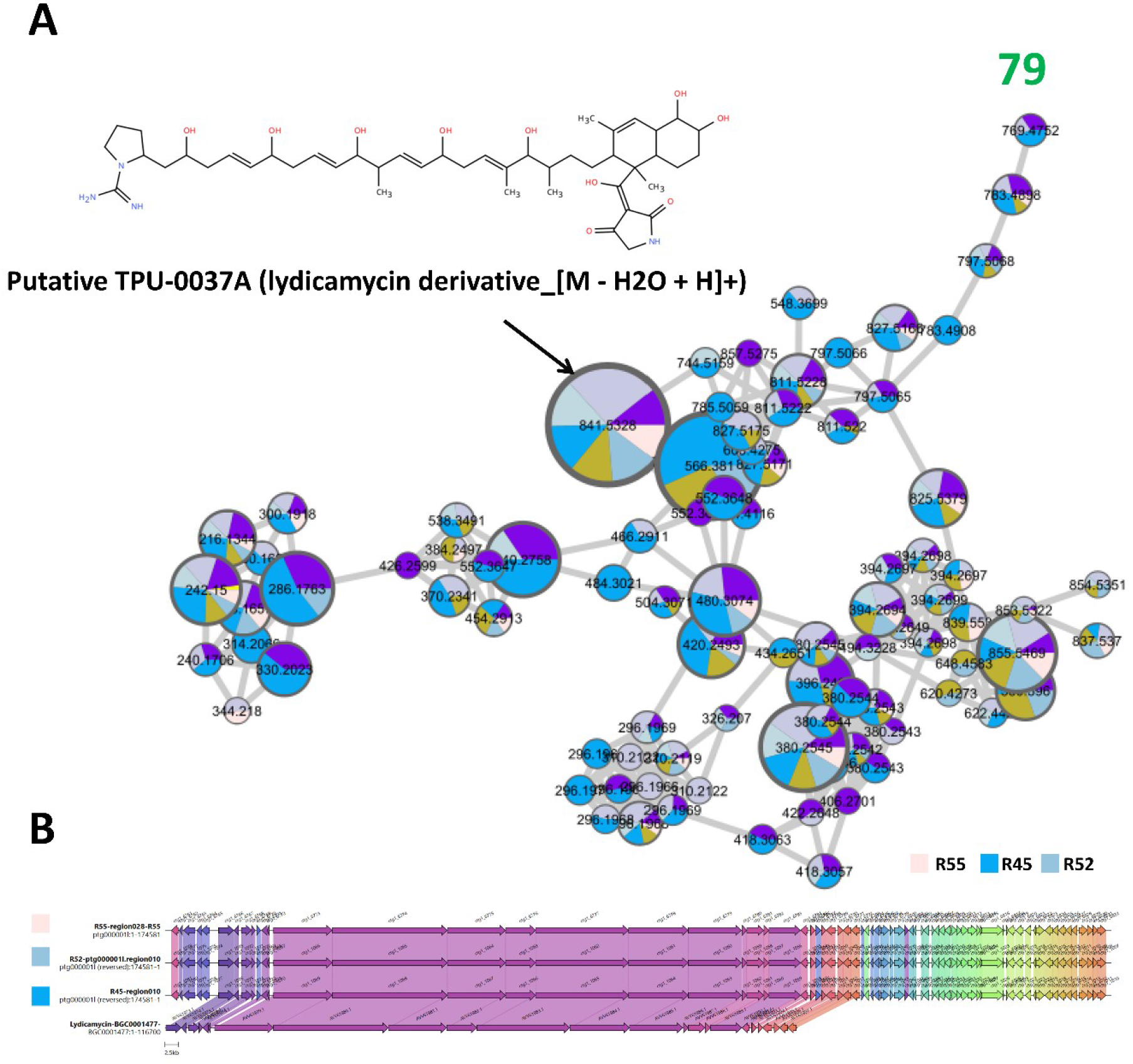
Genome-metabolome correlation of a putative lydicamycin-like molecular family. **(A)** FBMN-derived Component 79. Node colors indicate the distribution of metabolites across producing strains; individual strain labels were omitted to preserve figure readability due to the large number of contributing strains. **(B)** Comparison of lydicamycin-like T1PKS-NRPS BGCs identified in strains R45, R52, and R55 with the reference lydicamycin BGC (BGC0001477).

Comparative genome mining further reinforced this interpretation. Conserved lydicamycin-like T1PKS-NRPS biosynthetic gene clusters were identified in strains R45, R52, and R55, displaying strong synteny with the reference lydicamycin BGC (<u>BGC0001477</u>), particularly across the core T1PKS-NRPS region (Figure 10B). Together, the network organization, GNPS annotations, and mirror-spectrum comparison (Supplementary Figure S6) support the occurrence of a lydicamycin-related molecular family.

##### Tambromycin-associated molecular family (Component 44)

A peptide-rich molecular component (**Component 44**) was exclusively detected in strain R53 (Figure 11). The strongest evidence was provided by two central nodes <u>2473</u> (*m/z* 536.1915) and <u>2521</u> (*m/z* 536.1908), both independently annotated as <u>tambromycin A</u> by GNPS library matching (cosine scores 0.852 and 0.845, respectively). These central network nodes displayed fragmentation patterns consistent with the chlorinated tambromycin scaffold (Supplementary Figure S7). Several connected features, including <u>2472</u> (*m/z* 467.1332), <u>2520</u> (*m/z* 467.1327), <u>2535</u> (*m/z* 435.1434), and <u>2512</u> (*m/z* 796.3105), were classified as peptide- or depsipeptide-related metabolites by CANOPUS and ISDB-Lotus. Together, the GNPS annotations, peptide-related classifications, and network organization support the presence of a tambromycin-associated molecular family centered on tambromycin A, with additional unresolved congeners (Figure 11).

**Figure 11.**
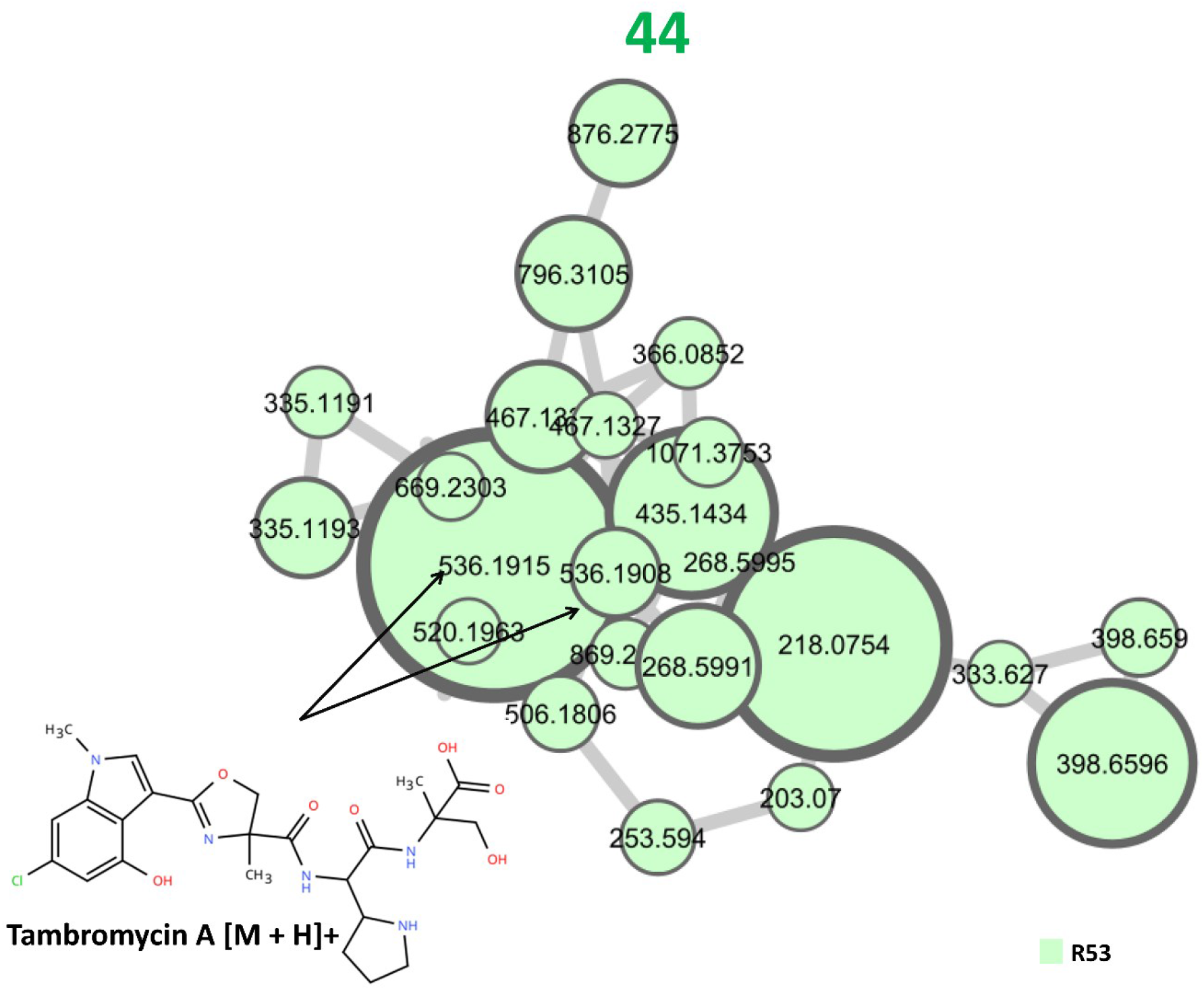
Molecular network of the tambromycin-associated molecular family detected exclusively in strain R53. Component 44 is centered on tambromycin A (features 2473 and 2521) and includes several connected peptide- and depsipeptide-related metabolites.

##### Hydroxamate siderophore-associated molecular family (Component 43)

**Component 43** corresponded to a molecular family enriched in hydroxamate siderophores and related polyamine-containing metabolites. The strongest GNPS annotations included <u>desferrioxamine G</u> (feature <u>1435</u>, *m/z* 619.3667), dehydroxynocardamine (feature <u>1704</u>, *m/z* 585.3600), <u>bisucaberin</u> (features <u>1260</u>, <u>1410</u>, and <u>1529</u>; *m/z* 401.2395-401.2398), and deferoxamine (feature <u>1422</u>, *m/z* 561.3612). Bisucaberin was detected as three closely related features exhibiting nearly identical masses and similar retention times (RT 2.34-2.66 min). Several additional connected nodes were classified by CANOPUS and ISDB-Lotus as polyamine-or hydroxamate-related metabolites, further supporting a siderophore-rich chemical space. The molecular family was primarily associated with strains R7B and R46C, while related features were also detected in other isolates, including R52 and R55. Collectively, the recurrent annotations and network topology support the presence of a diversified hydroxamate siderophore molecular family dominated by desferrioxamine-, nocardamine-, and bisucaberin-like metabolites.

Comparative genome mining further supported this interpretation. Strains R52 and R55 harbored NRPS-independent siderophore biosynthetic gene clusters displaying high similarity to the reference desferrioxamine B biosynthetic pathway (<u>BGC0000940</u>) (Figure 12B). Together, the metabolomic and genomic data support the production of hydroxamate siderophores within the strain collection.

**Figure 12.**
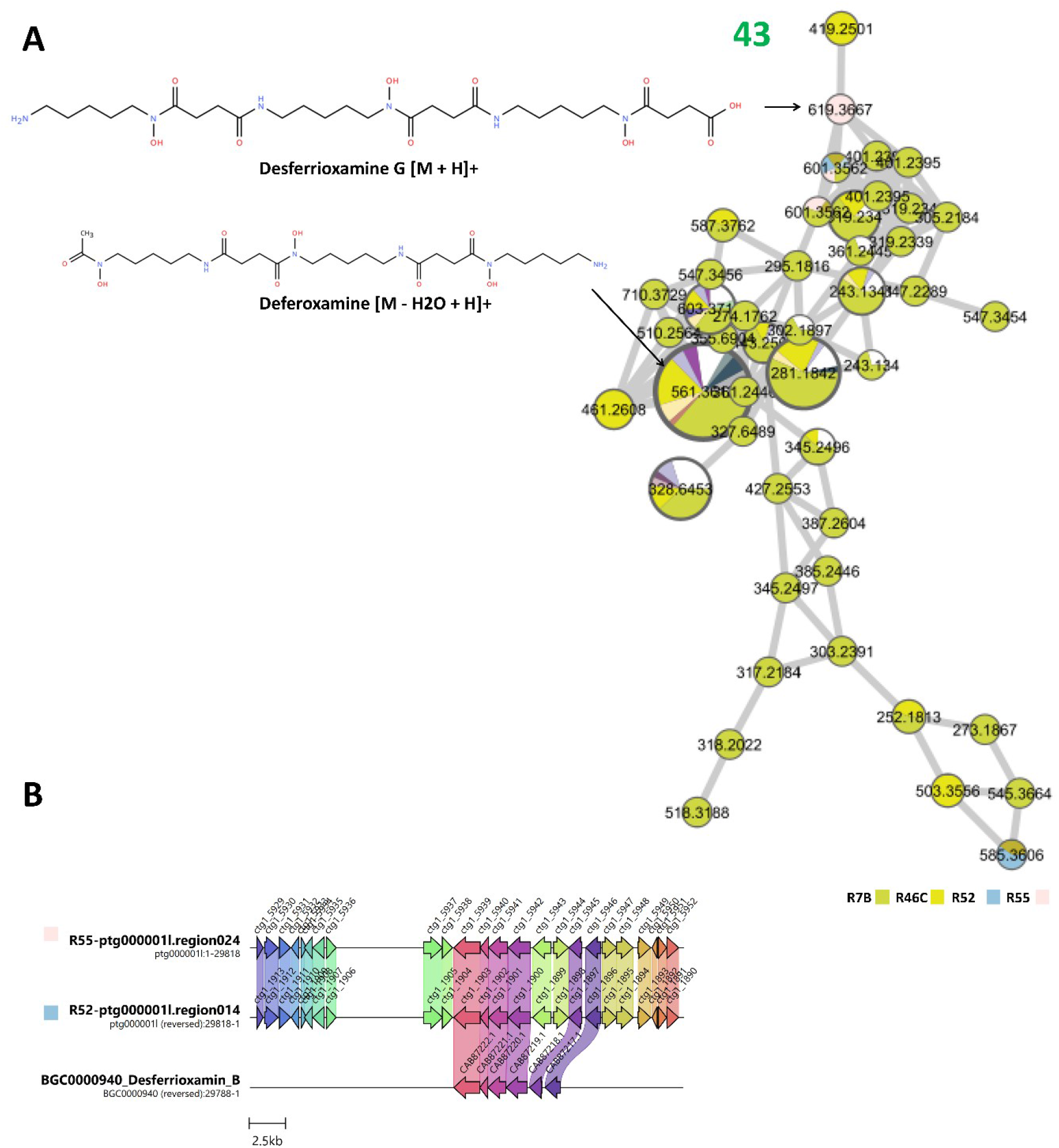
Genome-metabolome correlation of the hydroxamate siderophore-associated molecular family. **(A)** FBMN-derived Component 43. The displayed structures correspond to desferrioxamine G and deferoxamine. **(B)** Comparison of NRPS-independent siderophore BGCs identified in strains R52 and R55 with the reference desferrioxamine B BGC (BGC0000940).

##### Additional peptide- and amino acid-associated molecular families (Components 20, 182, 195, 229, 259, and 270)

Several molecular components were assigned to amino acid- and peptide-related chemical space (Figure 13A). CANOPUS classified the dominant nodes as amino acids and derivatives, dipeptides, tripeptides, oligopeptides, peptides, and peptidomimetics. Component 20 represented the largest peptide-rich molecular family (*m/z* 242.150-761.336), while Components 182, 195, 229, 259, and 270 comprised smaller peptide-enriched subnetworks. Collectively, these components define a diverse peptide-associated chemical space supported by FBMN, CANOPUS, SIRIUS, and ISDB-Lotus annotations, although no specific metabolite family could be confidently assigned.

Comparative genome mining identified several peptide-related BGCs in strains R54 and R56, including NRPS clusters showing 95% similarity to the Coprisamide C/D BGC (<u>BGC0002378</u>), cinnapeptin-like clusters (82% similarity), and a CDPS cluster in strain R54 displaying 100% similarity to the bicyclomycin BGC (Figure 13B). Despite this substantial peptide biosynthetic potential, no robust genome-to-metabolite correlations could be established.

**Figure 13.**
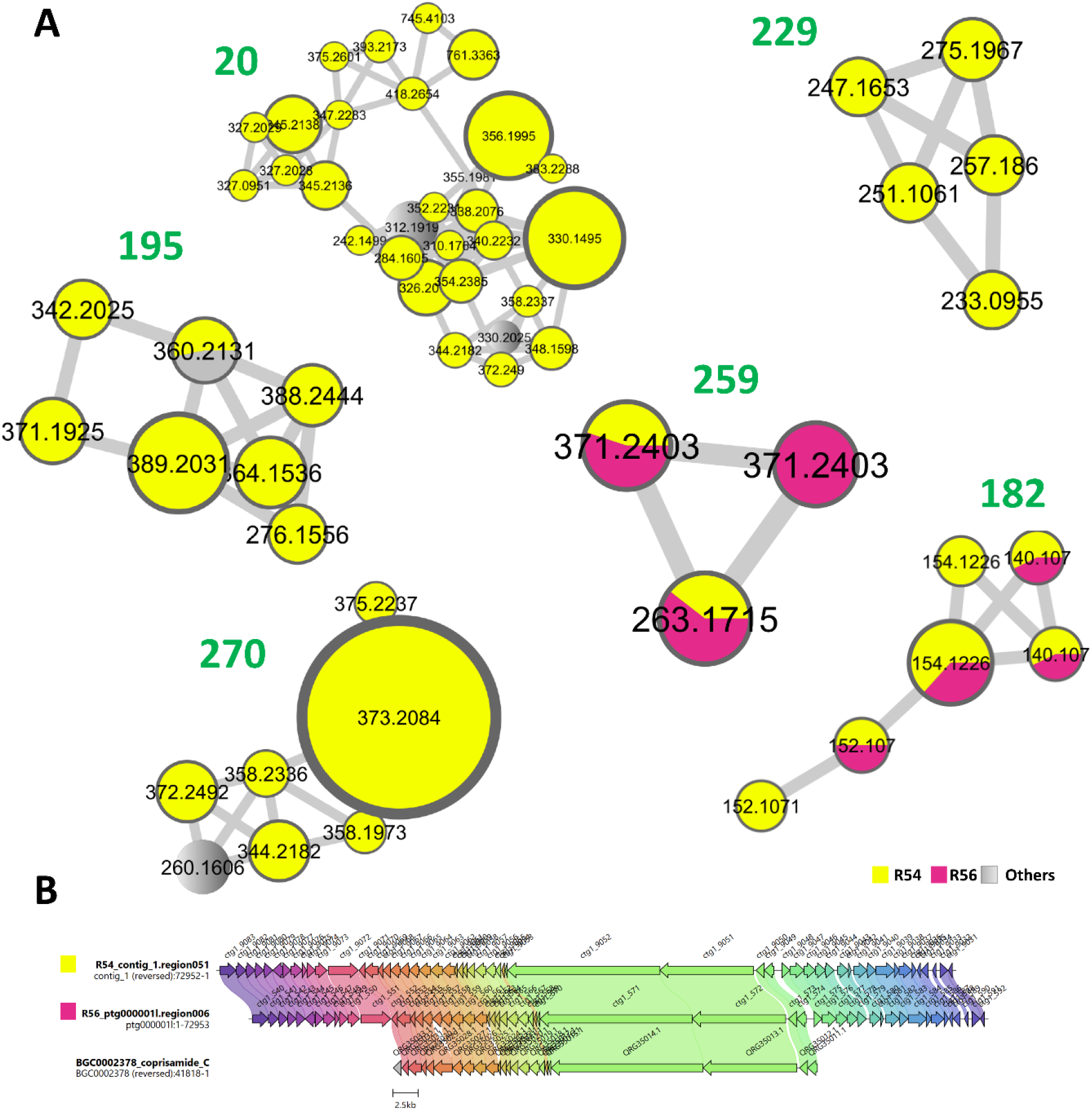
Metabolomic characterization and comparative genome mining of additional peptide-associated molecular families. **(A)** FBMN-derived Components 20, 182, 195, 229, 259, and 270. **(B)** Comparative alignment of Coprisamide-like NRPS BGCs from strains R54 and R56 with the reference Coprisamide C/D BGC (BGC0002378).

### 2.7 Operational assessment of the Duetz-based workflow

A proof-of-concept operational comparison was performed between the Duetz-based workflow and the conventional extraction procedure previously employed in our laboratory. In the conventional workflow, isolates were cultured in 200 mL of starch-casein broth, extracted with an equal volume of ethyl acetate, and processed into dried extracts prior to UHPLC-HRMS/MS analysis. In contrast, the Duetz workflow relied on cultivation in 1.8 mL of broth in 96-well Duetz microfermentation plates, followed by filtration through 0.22-µm 96-well filtration plates and direct UHPLC-HRMS/MS analysis.

For the 80-isolate collection investigated in this study, miniaturization reduced culture-medium consumption from 16 L to 0.144 L (99% reduction). Direct analysis of culture filtrates eliminated the use of 16 L of ethyl acetate and removed several sample-processing steps, reducing the workflow from eight to three principal preparation stages.

The plate-based format enabled parallel processing of up to 96 cultures and simplified sample handling through vacuum-assisted filtration. Based on laboratory experience, preparation and inoculation of an 80-isolate Duetz plate required approximately 30 min. Following cultivation, LC-MS-ready samples were obtained after a single ∼20 min plate-filtration step. In contrast, the conventional workflow typically required 6-8 h of active handling and 4-6 days of post-cultivation processing, including liquid-liquid extraction, phase separation, solvent evaporation, drying, and extract reconstitution, before LC-MS-ready extracts were obtained.

The Duetz workflow enabled antibacterial screening and metabolomic profiling from the same Duetz cultivation, thereby preserving a direct link between biological and chemical data. Recurring consumable costs decreased from approximately USD 401 to USD 16 per 80-isolate campaign (96.0% reduction), corresponding to a decrease from approximately USD 5.0 to USD 0.2 per isolate.

Overall, the Duetz-based workflow substantially reduced medium consumption, solvent use, processing time, and recurring consumable costs while enabling high-throughput, extraction-free metabolomic screening in a 96-well plate format. Table 1 summarizes the principal operational advantages of the workflow, whereas Supplementary Table S5 provides detailed material requirements, workflow steps, cost calculations, and processing-time estimates.

**Table 1.** Operational comparison of conventional and Duetz-based workflows for metabolomic screening.

| **Metric** | **Conventional workflow** | **Duetz-based workflow** | **Gain with the Duetz workflow** |
| --- | --- | --- | --- |
| **Culture format** | 80 individual flasks | One 96-well Duetz plate | Parallel cultivation |
| **Culture volume per isolate** | 200 mL | 1.8 mL | 111-fold reduction |
| **Total culture volume** | 16 L | 0.144 L | 99% reduction |
| **Ethyl acetate consumption** | 16 L | None | 100% elimination |
| **Sample preparation strategy** | Extraction, evaporation and reconstitution | Direct filtration and analysis | Extraction-free workflow |
| **Principal post-cultivation processing steps** | Eight stages | Three stages | Five stages eliminated (63% reduction) |
| **Processing mode** | Isolate-by-isolate | Plate-based and parallel | High-throughput processing of up to 96 samples simultaneously |
| **Time to obtain LC-MS-ready samples after cultivation** | Typically 4-6 days | ~20 min | ~360-fold reduction |
| **Workflow-specific recurring cost (80 isolates)** | USD 400 | USD 16 | USD 385 saved (96% reduction) |
| **Recurring cost per isolate** | USD 5 | USD 0.2 | USD 4.8 saved per isolate |
| **Bioactivity-metabolomics integration** | Separate workflows | Same Duetz cultivation used for both analyses | Direct integration of biological and chemical data |

## 3. Discussion

This study demonstrates the feasibility of integrating miniaturized cultivation, antibacterial screening, extraction-free metabolomics, and genome mining into a unified workflow for the rapid exploration of actinomycete collections. By combining these complementary approaches, we simultaneously assessed biological activity, metabolite production, and biosynthetic potential across a collection of 80 soil-derived actinomycetes. This multi-omics framework enabled efficient strain prioritization while providing a comprehensive overview of the chemical and genomic diversity present within the collection.

A major advantage of the approach lies in its operational efficiency, scalability, and cost-effectiveness. Compared with our previous extraction-based strategy, the Duetz workflow reduced culture-medium consumption by 99%, eliminated the use of 16 L of ethyl acetate during an 80-isolate screening campaign, and decreased recurring consumable costs by approximately 96%. Direct analysis of culture filtrates removed several labor-intensive processing steps, including liquid-liquid extraction, solvent evaporation, drying, and extract reconstitution, substantially reducing sample handling and preparation time. The 96-well Duetz format further enabled parallel cultivation and processing of large microbial strain collections, highlighting the value of workflow miniaturization for increasing screening capacity while reducing material consumption, processing time, and operational costs.

Beyond these practical advantages, the integrated workflow provided a comprehensive view of strain bioactivity, chemistry, and biosynthetic potential within a single experimental framework. Although compound isolation and structure elucidation remain essential for definitive characterization, the approach facilitates informed decision-making early in the discovery process and can be readily applied to large microbial collections. By integrating biological, metabolomic, and genomic information, the workflow supports robust strain prioritization and the identification of biologically relevant metabolites for downstream investigation.

Seven isolates displayed antagonistic activity against Gram-positive pathogens, including *S. aureus* and *E. faecium*, further supporting the recognized potential of actinomycetes as a source of bioactive microorganisms.^24–26^ Activity was restricted to methicillin-resistant *Staphylococcus aureus* (MRSA) and *E. faecium*, whereas no inhibition was detected against the Gram-negative pathogens tested. This pattern is consistent with the intrinsic permeability barrier and multidrug-resistance mechanisms commonly associated with Gram-negative bacteria, which reduce susceptibility to many natural products.^27^ The relatively small proportion of active isolates likely reflects the condition-dependent nature of specialized metabolite production in actinomycetes, whereby only a subset of biosynthetic pathways is expressed under a given cultivation condition. Medium composition, nutrient availability, aeration, and other cultivation parameters are known to strongly influence metabolite production and associated biological activities.^28,29^

Importantly, genome mining revealed that several inactive isolates nevertheless harbored diverse biosynthetic gene clusters, highlighting the frequent disconnect between biosynthetic potential and metabolite expression under a single cultivation condition. This finding underscores the value of integrating antibacterial screening with metabolomic and genomic analyses to identify promising strains that would otherwise be overlooked.

The taxonomic structure of the sequenced subset revealed a marked predominance of *Streptomyces*, consistent with the widespread occurrence of this genus in soil environments and its frequent recovery using conventional actinomycete isolation procedures.^30–32^ Taxonomic assignments based on GTDB-Tk and ANI analyses were highly consistent. Several isolates displayed ANI values close to species delineation thresholds^33^, suggesting the presence of genetically differentiated lineages within the collection. This diversity is particularly relevant for natural product discovery because biosynthetic gene cluster content can vary even among closely related strains.^34,35^ Consistent with this observation, a broad diversity of biosynthetic gene clusters and metabolite families was identified across the collection, highlighting the extensive metabolic and biosynthetic potential of the investigated actinomycetes.

The inclusion of inactive strains provided a more comprehensive view of metabolomic and genomic diversity across the collection. Genome-metabolome integration revealed molecular families associated with antibacterial-active isolates, including actinomycins, polyether ionophores, and ansamycins, whereas angucycline-, ectoine-, and lydicamycin-associated families were predominantly detected in inactive isolates. Additional peptide-rich molecular families and hydroxamate siderophores were distributed across multiple strains and reflected broader ecological and physiological functions. The level of agreement between molecular networking, metabolite annotation, GNPS library matching, and biosynthetic gene cluster prediction varied across metabolite families, resulting in different levels of confidence in the proposed genome-metabolome associations. Nevertheless, this integrative analysis enabled the identification of both bioactivity-associated and ecologically relevant molecular families, highlighting the value of including inactive strains in discovery-oriented screening workflows. Among the prioritized families, the actinomycin, polyether ionophore, and ansamycin families displayed the strongest agreement between metabolomic and genomic datasets. In each case, metabolite annotations were supported by biosynthetic gene clusters showing high similarity to experimentally characterized pathways, increasing confidence in the proposed genome-metabolome associations and metabolite annotations. Nevertheless, although several key metabolites, including actinomycin D, geldanamycin, ectoine, 5-hydroxyectoine, monensin A, and nigericin, were further supported through comparison with commercial standards, many compounds remain assigned at the molecular-family level and await further structural confirmation.

The actinomycin family may contribute to the antibacterial activity observed in the most active isolates. Actinomycins are among the best-characterized antibiotics produced by *Streptomyces* and are known for their potent activity against Gram-positive bacteria through DNA intercalation and inhibition of transcription.^36,37^ Their predominance among the most active strains, combined with the agreement between metabolomic and genomic data and confirmation of actinomycin D using a commercial standard, supports their involvement in the antibacterial activity observed in this study.

The polyether ionophore family was associated with isolates displaying antibacterial activity against Gram-positive pathogens. Members of this metabolite class are well known for their ability to disrupt ion homeostasis by acting as mobile cation transporters across biological membranes, resulting in potent activity against Gram-positive bacteria.^38,39^ Their detection in strains R54, R56, and R60 is therefore consistent with the antibacterial profiles observed in this study. Notably, all three strains were assigned to *Streptomyces hygroscopicus* (Supplementary Table S1, Sheet 2), a species known to produce structurally diverse polyether ionophores. The occurrence of multiple related congeners together with highly similar biosynthetic gene clusters, and the confirmation of monensin A and nigericin using commercial standards, further strengthens the proposed genome-metabolome association. Nevertheless, the relative contribution of individual ionophores to the observed antibacterial activity remains to be determined.

The ansamycin molecular family detected in strains R54 and R56 further illustrates the complementarity between metabolomic and genomic datasets. The occurrence of geldanamycin-, macbecin-, and herbimycin-related metabolites, together with a herbimycin-like biosynthetic gene cluster, supports the assignment of this molecular family to benzoquinone ansamycins. Members of this group are known for diverse biological activities, notably through inhibition of the molecular chaperone Hsp90.^40,41^ At the same time, the strong structural similarity shared by many ansamycins highlights the limitations of metabolite dereplication based solely on MS/MS data. Both R54 and R56 were assigned to *Streptomyces hygroscopicus* (Supplementary Table S1, Sheet 2), a species known to produce herbimycin-type ansamycins.^42^ In this context, the convergence of molecular networking, metabolite annotation, taxonomic information, and genome mining enabled assignment of these metabolites to a common biosynthetic family while avoiding overinterpretation at the individual compound level.

In contrast to the metabolite families associated with antibacterial activity, the angucycline-associated molecular family was predominantly detected in inactive isolates and comprised fridamycin and related angucycline metabolites. Angucyclines represent a large group of aromatic type II polyketides produced by *Streptomyces* and are known to display diverse biological activities.^43^ In this study, they were primarily detected in strains R46C and R7A, with R46C representing the major producer. Their occurrence in inactive isolates highlights the complexity of linking metabolite production to biological activity, as the presence of bioactive metabolites does not necessarily translate into measurable antibacterial activity under a given screening condition. These findings illustrate how metabolomic profiling can uncover chemically diverse and potentially bioactive metabolites that may be overlooked through activity-guided prioritization alone.

The putative lydicamycin-like molecular family represents an interesting example of the complementary insights provided by metabolomic and genomic analyses. Although not directly associated with detectable antibacterial activity under the tested conditions, lydicamycin-related metabolites have previously been reported to display potent activity against Gram-positive pathogens.^44^ The detection of a lydicamycin-like molecular family together with conserved T1PKS-NRPS biosynthetic gene clusters provides complementary metabolomic and genomic evidence supporting the occurrence of lydicamycin-related metabolites within the collection.

Beyond metabolite families potentially linked to antibacterial activity, Component 43 highlighted the widespread distribution of hydroxamate siderophores across the collection. Metabolites annotated as desferrioxamine-, bisucaberin-, and nocardamine-related compounds are involved in microbial iron acquisition under iron-limited conditions^45,46^. Their occurrence across multiple isolates, together with the identification of desferrioxamine-like biosynthetic gene clusters in strains R52 and R55, supports the widespread distribution of hydroxamate siderophore biosynthesis within the collection.

The ectoine-associated family was predominantly detected in inactive isolates and appears to reflect physiological adaptation rather than direct antibacterial activity. Ectoine and 5-hydroxyectoine are widely distributed compatible solutes that contribute to osmotic balance and stress tolerance in diverse bacteria, including actinomycetes.^47,48^ The detection of ectoine-associated metabolites together with conserved ectoine biosynthetic gene clusters across multiple isolates provides concordant metabolomic and genomic evidence for ectoine biosynthesis within the collection. Their predominance in inactive isolates is therefore consistent with a role in environmental adaptation and ecological fitness rather than antagonistic interactions.

Together, the lydicamycin-, angucycline-, ectoine-, and hydroxamate siderophore-associated families illustrate how metabolomic and genomic analyses can reveal biologically and ecologically relevant traits that would likely remain undetected through activity-guided screening alone.

In contrast to molecular families distributed across multiple isolates, Component 44 was detected exclusively in strain R53 and was centered on tambromycin A. Tambromycins constitute a family of halogenated nonribosomal peptides produced by actinomycetes and have been reported to exhibit antiproliferative activity.^49^ The detection of tambromycin A highlights the unique biosynthetic potential of R53. Although no genome sequence is currently available for this isolate, preventing direct genome-metabolome correlation, the occurrence of this molecular family underscores the value of metabolomic screening for identifying chemically unique producers and makes R53 an attractive target for future genome sequencing and chemical characterization.

Several additional peptide-rich molecular families **(Components 20, 182, 195, 229, 259**, and **270**) were identified across the collection. Although their occurrence was consistent with the presence of peptide-associated NRPS and CDPS biosynthetic systems in strains R54 and R56, the absence of direct genome-to-metabolite evidence precluded confident assignment to specific biosynthetic pathways. These molecular families therefore mainly reflect the broader peptide biosynthetic potential of the investigated strains.

While the integrated workflow enabled the prioritization of bioactive isolates and the identification of metabolite families of ecological and biosynthetic interest, several limitations should be considered when interpreting these data. First, the metabolomic analysis was restricted to metabolites present in cell-free culture filtrates and may therefore underrepresent intracellular, cell-associated, or highly hydrophobic compounds. Second, all strains were analyzed under a single cultivation condition, although the expression of many biosynthetic pathways is strongly influenced by environmental and nutritional factors.^50,51^ This may explain why certain molecular families previously associated with biological activity, such as the lydicamycin- and angucycline-associated families, were predominantly detected in isolates that did not exhibit antibacterial activity under the tested conditions. Furthermore, most metabolite annotations reported in this study should be regarded as putative family-level assignments based on MS/MS spectral similarity, in silico annotation, and genome mining, and will ultimately require validation through compound isolation and orthogonal structural characterization.

Despite these limitations, the Duetz microfermentation format is particularly well suited to OSMAC (One Strain Many Compounds) strategies, ^52,29^ as strains can be readily replicated from a common stock collection into multiple 96-well cultivation plates containing different media or growth conditions. Consequently, the workflow provides a practical framework for large-scale OSMAC experiments aimed at activating silent or poorly expressed pathways and expanding the accessible chemical space of actinomycete collections.

Overall, this study demonstrates how miniaturized cultivation can support natural product discovery through the integrated generation of bioactivity, metabolomic, and genomic data at collection scale. The proposed workflow enabled the prioritization of both antibacterial-active isolates and strains harboring metabolite families of ecological or biosynthetic interest, including angucycline-, lydicamycin-, ectoine-, hydroxamate siderophore-, and tambromycin-associated metabolites, as well as several peptide-rich molecular families. Despite the absence of solvent-based extraction during primary screening, direct analysis of culture filtrates enabled the annotation of diverse metabolite families and the establishment of biologically meaningful genome-metabolome associations across the collection. These findings demonstrate that culture filtrates can provide sufficient chemical information for strain selection while substantially simplifying sample processing. The flexibility, low material requirements, and scalability of the Duetz platform make it particularly well suited for future OSMAC-based discovery campaigns, in which strains can be systematically evaluated under multiple cultivation conditions in parallel. Combined with advances in metabolomics, genome mining, and computational annotation, such approaches have the potential to improve the identification of microorganisms harboring both expressed and cryptic biosynthetic potential while reducing the time, cost, and material requirements associated with large-scale discovery efforts. More broadly, this study highlights the value of integrating biological, chemical, and genomic information to accelerate the exploration of microbial specialized metabolism and support informed strain prioritization for natural product discovery.

## 4. Experimental section

### 4.1 Sampling and pretreatment of soil

Soil samples were collected from diverse terrestrial biotopes across Switzerland (Supplementary Table S1, Sheet 1) and transported to the laboratory in sterile Falcon tubes. To reduce the recovery of fast-growing microorganisms while promoting the isolation of rare actinomycete genera, each sample underwent a pretreatment step before cultivation. For heat pretreatment, approximately 1 g of dry soil was exposed to dry heat at 100 °C for 1 h in a hot-air oven, as previously described,^53^ (T1, Supplementary Table S1). For chemical pretreatment, 1 g of soil was suspended in 10 mL of sterile NaCl solution (0.9%) and treated with 1.5% phenol for 30 min at 30 °C, according to^54^ (T2, Supplementary Table S1). Following pretreatment, soil samples were suspended in sterile NaCl solution (0.9%), and 100 µL aliquots were spread onto Starch-Casein Agar (SCA) supplemented with nalidixic acid (10 µg/mL), nystatin (50 µg/mL), and cycloheximide (50 µg/mL). Three replicate plates were prepared for each pretreatment and sample. Plates were incubated at 28 °C for up to 3 months. After incubation, colonies displaying morphological characteristics typical of actinomycetes were selected and purified by successive streaking on fresh SCA plates. To maximize isolate diversity, colonies displaying distinct morphotypes were preferentially selected based on pigmentation, colony texture, growth pattern, and aerial mycelium formation. This strategy was used solely to reduce the recovery of redundant isolates and increase the likelihood of capturing phenotypic diversity within the collection. Morphological characteristics were not used for taxonomic identification or to infer phylogenetic relationships among isolates.

### 4.2 Assessment of antibacterial activity against ESKAPE pathogens

A total of 80 purified actinomycete isolates were included in the antibacterial screening. To minimize inoculum variability, each culture was initiated using a standardized 5 mm agar plug excised from an actively growing colony on SCA medium. Isolates were cultivated in 1.8 mL of starch-casein broth in 96-well Duetz microfermentation plates and incubated at 28 °C for 14 days under orbital shaking (110 rpm) to promote specialized metabolite production.

Antibacterial activity was assessed against *Staphylococcus aureus* ATCC 43300, *Enterococcus faecium* BM4147, *Klebsiella pneumoniae* ATCC 700603, *Escherichia coli* ATCC 25922, *Acinetobacter baumannii* CIP 7010, and *Pseudomonas aeruginosa* ATCC 27853. For the antagonism assay, Mueller-Hinton agar plates were first inoculated with each pathogen at a final concentration of 6 × 10⁵ CFU/mL.^55^ Subsequently, 4 µL of each actinomycete culture broth was directly spotted onto the pathogen-seeded agar surface. Plates were then maintained at room temperature for 2 h to facilitate metabolite diffusion prior to incubation at 37 °C for 24 h. Following incubation, inhibition zone diameters were measured. Strains producing visible inhibition zones were classified as active. Metabolomic analyses were performed on all isolates, whereas active strains were prioritized for further investigation and genome sequencing.

### 4.3 Genome sequencing, assembly, and taxonomic assignment

Each actinomycete strain was grown in 40 mL of Tryptic Soy Broth (TSB; Sigma-Aldrich) at 28 °C under shaking conditions (180 rpm) for 3 days. Genomic DNA was extracted using the GenElute™ Bacterial Genomic DNA Kit (Merck) following the manufacturer’s protocol, with extended incubation times for lysozyme (90 min instead of 30 min) and proteinase K/lysis solution C (30 min instead of 10 min). DNA quality was assessed using 1% agarose gel electrophoresis, and concentration was determined with a NanoDrop spectrophotometer. Genomic DNA was sequenced at the University of Bern Next-Generation Sequencing Platform using Pacific Biosciences (PacBio) library preparation protocols. Libraries were multiplexed on a single SMRT Cell 25M and sequenced on the PacBio Revio system to generate HiFi long reads. Demultiplexing was performed with lima, and read quality was assessed with FastQC.

Initial assemblies were generated using the SMRT Link pb_microbial_analysis pipeline (v11.0.0.146107), which also provided polishing, circularization, and origin rotation when required. The pipeline also identified DNA modifications and methylated motifs. Genome completeness and contamination were evaluated using CheckM2.^56^

Following quality assessment and filtering, de novo assemblies were generated using the SMRT Link Microbial Assembly pipeline (Pacific Biosciences), Hifiasm^57^, and Flye.^58^ The resulting assemblies were compared based on completeness, contamination, contiguity metrics, and circularization status when applicable. The highest-quality assembly was retained for downstream analyses. Taxonomic classification was performed using Kraken2^59^ and Bracken^60^ on raw sequencing reads and GTDB-Tk^61^ on assembled genomes, and Average Nucleotide Identity (ANI) analyses were performed for comparative genomic assessment of strain relationships.

A total of 39 strains were selected for long-read genome sequencing, including four of the seven isolates displaying antibacterial activity. The three remaining active isolates were not sequenced because insufficient amounts of high-quality genomic DNA were obtained for library preparation. The remaining strains were selected to capture distinct colony morphotypes and maximize the phenotypic diversity of the collection across sampling sites. Inactive isolates were intentionally included because the absence of detectable antibacterial activity under a single cultivation condition does not necessarily indicate a lack of biosynthetic potential, as many BGCs remain poorly expressed or silent under laboratory conditions.

### 4.4 Genome mining of biosynthetic gene clusters

The relaxed detection setting was selected to maximize BGC recovery during the exploratory phase of strain prioritization. Predicted BGCs were subsequently examined and compared based on their similarity to experimentally characterized reference clusters available in the antiSMASH^62^ and MIBiG database^63^. Similarity values reported by antiSMASH were used to assess relationships between predicted BGCs and previously characterized biosynthetic pathways. BLASTp analyses against the NCBI non-redundant protein database were performed when necessary to support cluster annotation and refine the functional assignment of selected biosynthetic genes.

### 4.5 Cultivation and sample preparation for UHPLC-HRMS Analysis

The actinomycete collection was cultivated in 96-well Duetz microfermentation plates containing 1.8 mL of starch-casein broth per well and incubated at 28 °C for 14 days under orbital shaking at 110 rpm. Following incubation, 1.8 mL of each culture was transferred to a 96-well filter plate equipped with a 0.22 µm membrane and placed on a deep-well collection plate. Cell-free culture filtrates were obtained by vacuum-assisted filtration, which retained bacterial biomass on the membrane while allowing extracellular metabolites to pass into the collection plate.

Unlike conventional natural product discovery workflows, no solvent-extraction step was performed prior to UHPLC-HRMS analysis. This approach enabled direct metabolomic profiling of cell-free culture filtrates while supporting high-throughput screening and reducing sample-handling requirements and processing-related variability. The resulting filtrates were analyzed directly by ultra-high-performance liquid chromatography coupled to high-resolution mass spectrometry (UHPLC-HRMS).

A schematic overview of the Duetz-based cultivation, filtration, and extraction-free UHPLC-HRMS workflow used for metabolomic analysis is presented in Figure 14.

**Figure 14.**
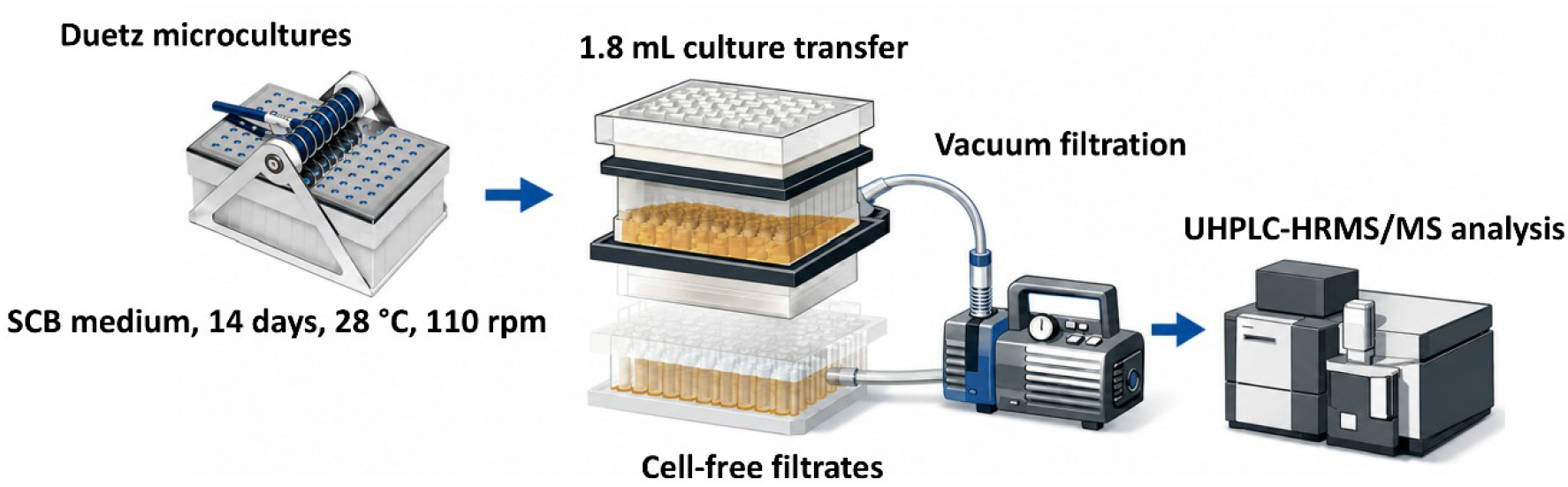
Overview of the Duetz-based cultivation, filtration, and sample-preparation workflow for direct UHPLC-HRMS/MS analysis.

### 4.6 UHPLC-HRMS profiling of cell-free filtrates from the actinomycete collection

The previously prepared cell-free culture filtrates were subjected to untargeted metabolomic analysis. Metabolite profiling was performed using reverse-phase chromatography coupled with electrospray ionization and accurate-mass detection. Chromatographic separation was carried out on a Vanquish Flex UPLC system (Thermo Fisher Scientific) interfaced with a Q-Exactive HF-X mass spectrometer (Thermo Fisher Scientific) equipped with a heated electrospray ionization (HESI-II) source. Instrument control was performed using Thermo Scientific Xcalibur 3.1 software. Culture-medium blanks were included throughout the workflow and used to identify medium-derived signals during molecular networking analyses. Features predominantly associated with medium-control samples were considered background signals and were excluded from biological interpretation. Quality control (QC) samples were analyzed at regular intervals throughout the UHPLC-HRMS acquisition sequence to monitor retention time stability, mass accuracy, analytical reproducibility, and overall instrument performance.

LC conditions were as follows: Waters BEH C18 column (100 × 2.1 mm, 1.7 μm); mobile phase A, water containing 0.1% formic acid; mobile phase B, acetonitrile containing 0.1% formic acid; flow rate, 600 μL/min; injection volume, 2 μL; linear gradient from 2% to 100% B over 10 min, followed by an isocratic step at 100% B for 2 min.

The optimized HESI-II parameters were: source voltage, 3.5 kV (positive mode); sheath gas flow rate (N₂), 55 units; auxiliary gas flow rate, 15 units; sweep gas flow rate, 3 units; capillary temperature, 350 °C; and S-Lens RF level, 50.

The mass spectrometer was calibrated by direct infusion of a standard mixture containing caffeine, methionine-arginine-phenylalanine-alanine, acetate, sodium dodecyl sulfate, sodium taurocholate, and Ultramark 1621 dissolved in acetonitrile/methanol/water containing 1% formic acid.

Full-scan MS data were acquired in positive ionization mode over an *m/z* range of 140–2000. Data-dependent MS/MS acquisition was performed on the three most intense precursor ions detected in each full-scan MS spectrum (Top3 method). The isolation window was set to 1 Da, and stepped normalized collision energies of 15, 30, and 45 were applied. Full-scan MS spectra were acquired at a resolution of 35,000 FWHM (*m/z* 200), whereas MS/MS spectra were acquired at 17,500 FWHM, both using automatically determined maximum injection times. Following MS/MS acquisition, precursor ions were excluded from further fragmentation for 3 s using dynamic exclusion.

### 4.7 Data processing and molecular networking

Raw UHPLC-HRMS/MS data were converted to mzML format using ProteoWizard MSConvert and processed with the MZmine software package (v4.5.0).^64^ Mass detection was performed using the centroid mass detector, with the noise level set to 5.0 × 10⁶ for MS level 1 and 0 for MS/MS scans. Chromatograms were built using the ADAP Chromatogram Builder with a minimum group size of 5 scans, a minimum intensity threshold of 5.0 × 10⁶, a minimum highest intensity of 5.0 × 10⁶, and an *m/z* tolerance of 15 ppm. Chromatograms were subsequently smoothed using the Savitzky-Golay algorithm with a retention time smoothing window of 5 scans. Chromatogram deconvolution was performed using the Local Minimum Search algorithm with a chromatographic threshold of 0.3, a minimum search range of 0.05 min, a minimum feature height of 5.0 × 10⁴, a minimum peak-top-to-edge ratio of 1.8, a peak duration range of 0-1.51 min, and a minimum of 5 data points per peak. MS/MS spectra were associated with chromatographic features using a precursor *m/z* tolerance of 10 ppm and a retention time tolerance of 0.05 min. Isotopes were grouped using the Isotope Grouper module with an *m/z* tolerance of 5 ppm, a retention time tolerance of 0.04 min, and a maximum charge state of 2, using the most intense isotope as the representative peak. Peak alignment was performed using the Join Aligner algorithm with an *m/z* tolerance of 5 ppm and a retention time tolerance of 0.1 min, with weights of 3 and 2 assigned to *m/z* and retention time, respectively. Feature filtering retained only features associated with MS/MS spectra and containing at least two isotopic peaks. The final aligned feature list, including the feature quantification table and associated MS/MS spectra, was exported for GNPS Feature-Based Molecular Networking (FBMN) and SIRIUS analyses.

Molecular networking was performed on the GNPS2 platform,^65^ using the exported .mgf file. The precursor ion mass tolerance was set to 0.02 Da, and an MS/MS fragment ion tolerance of 0.02 Da. A network was then created where edges were filtered to have a cosine score above 0.7 and more than six matched peaks. Further, edges between two nodes were kept in the network if and only if each node appeared among the other’s top 10 most similar nodes. Finally, the maximum size of a spectral family was set to 100, and the lowest-scoring edges were removed from molecular families until the molecular family size was below this threshold. The spectra in the network were then searched against GNPS spectral libraries. All matches kept between network spectra and library spectra were required to have a score above 0.7 and at least six matched peaks.

### 4.8 Metabolite annotation strategy

Metabolite annotation was performed using complementary experimental and in silico approaches. First, MS/MS spectra were compared against the GNPS2 experimental spectral libraries to identify metabolites supported by reference spectra. Second, annotations were refined through the integration of spectral matching, taxonomic information, and in silico approaches. A theoretical natural products (NP) spectral database was generated according to a previously described workflow.^66,67^ The information used to build this resource originates from the LOTUS electronic resources hosted on Wikidata, which compile natural products and their biological occurrences into an open and continuously evolving database.^68,69^ This database, together with associated biological source metadata, is publicly available.^70^ Taxonomically informed metabolite annotation was performed using the met_annot_enhancer scripts (v0.1) available on GitHub: https://github.com/mandelbrot-project/met_annot_enhancer/releases/tag/v0.1. In addition, SIRIUS (v6.3.0),^71^ including CSI: FingerID, was used for molecular formula prediction and structural annotation, whereas CANOPUS^72,73^ was used for chemical class prediction from MS/MS spectra. Metabolite annotations were interpreted according to the combined evidence provided by GNPS library matching, taxonomically informed annotation, SIRIUS molecular formula prediction, CANOPUS chemical classification, molecular networking, and genome-mining analyses.

### 4.9 Validation of selected metabolite annotations using commercial standards

Commercial standards were prepared according to the manufacturers’ recommendations and analyzed individually using the chromatographic and mass spectrometric conditions described above. Retention times, accurate masses, molecular formulas, and ion species were compared with the corresponding LC-MS features detected in culture filtrates. When metabolite signals were weak or not readily detectable in pooled QC samples due to dilution effects, comparisons were additionally performed using individual culture filtrates. Representative chromatographic comparisons are provided in Supplementary Figure S8, whereas detailed agreements in retention time, accurate mass, molecular formula, and adduct type are summarized in Supplementary Table S6.

## 5. Conclusions

This study highlights the potential of integrating miniaturized cultivation, antibacterial screening, extraction-free metabolomics, and genome mining into a unified workflow for the exploration of actinomycete chemical diversity. The combination of these complementary approaches enabled the identification of antibacterial strains, the characterization of 24 prioritized molecular components spanning diverse specialized metabolite families, and the establishment of multiple putative genome-metabolome associations supported by biosynthetic gene cluster analyses.

A key strength of the workflow lies in its ability to generate biological, chemical, and genomic information from the same miniaturized cultivation platform. Direct UHPLC-HRMS analysis of cell-free culture filtrates eliminated extraction-related processing during primary screening while reducing material consumption, sample handling, and operational costs. Despite the absence of solvent extraction, the approach provided sufficient chemical information to support metabolite dereplication, molecular family annotation, and genome-metabolome investigations. Confidence in selected annotations was further strengthened through comparison with commercial standards.

Although no previously undescribed metabolite families were confidently identified under the cultivation conditions tested and antibacterial activity was restricted to Gram-positive pathogens, the inclusion of both active and inactive isolates revealed molecular families associated with antibacterial activity, ecological adaptation, and cryptic biosynthetic potential. The scalability and flexibility of the Duetz platform make it particularly well suited for future OSMAC-based investigations and large-scale screening campaigns. Combined with advances in metabolomics, genome mining, and computational annotation, this strategy provides a practical framework for accelerating microbial natural product discovery while reducing the time, cost, and material requirements associated with large-scale strain exploration.

## ASSOCIATED CONTENT

### Supporting Information

Supplementary **Table S1**. Metadata associated with the actinomycete collection, including isolate information, sampling data, pretreatment conditions, taxonomic assignments, and genome accession numbers. Supplementary **Table S2.** Antibacterial screening results, including inhibition zone diameters of actinomycete isolates against the tested bacterial pathogens. Supplementary **Table S3.** Biosynthetic gene clusters predicted by antiSMASH for the 39 sequenced genomes. Supplementary **Table S4.** Molecular feature annotations obtained from feature-based molecular networking (FBMN), including feature IDs, molecular component assignments, and annotations generated using GNPS, ISDB-Lotus, SIRIUS, and CANOPUS. **Supplementary Table S5.** Operational comparison of conventional and Duetz-based workflows. Supplementary **Table S6**. Comparison of selected metabolite annotations with commercial standards, including retention time, accurate mass, molecular formula, and observed ion species. **Supplementary Figures.** GNPS mirror-spectrum validations supporting the putative annotation of selected metabolite families (**Figures S1-S7**) and extracted ion chromatogram (EIC) comparisons between commercial standards and corresponding LC-MS features supporting selected metabolite annotations (**Figure S8**).

### Accession codes

The genomic data generated in this study have been deposited in the European Nucleotide Archive (ENA) under project accession number **PRJEB108104**. The corresponding study accession is **ERA35660740**. The data are available through the ENA browser. Detailed accession numbers for the project, study, samples, experiments, and sequencing runs are provided in Supplementary Table S1 (Sheet 3). The complete LC-MS/MS dataset, FBMN results, and associated metadata have been deposited in the MassIVE repository under accession **MSV000103110** and are publicly available through the associated dataset DOI: https://doi.org/10.25345/C5XP6VJ31. The corresponding GNPS2 Feature-Based Molecular Networking (FBMN) job is publicly available at: https://gnps2.org/status?task=a2721f67b9af435da7be0db18b25fbae.

### NOTES

The authors declare no competing financial interests.

## AUTHOR CONTRIBUTIONS

### Author contributions were defined according to the <u>CRediT</u> taxonomy

**RO:** Conceptualization, Methodology, Investigation, Formal Analysis, Validation, Data Curation, Visualization, Writing – Original Draft, Writing – Review & Editing.

**LF:** Formal Analysis, Data Curation, Writing – Review & Editing.

**LP:** Resources, Writing – Review & Editing.

**PMA:** Conceptualization, Methodology, Formal Analysis, Data Curation, Resources, Supervision, Project Administration, Funding Acquisition, Writing – Review & Editing.

## FUNDING

This work was supported by the Swiss Confederation through a Swiss Government Excellence Scholarship awarded to Dr. Rima Ouchene and by partial funding from the Centenary Research Fund of the University of Fribourg.

## ACKNOWLEDGMENTS

The authors thank Prof. Laure Weisskopf for access to laboratory facilities and Dr. Floriane L’Haridon, Dr. Ola Abdelrahman, and Mr. Maxime Bouvier for technical assistance.

## ABBREVIATIONS

ANI: Average Nucleotide Identity
BGC: Biosynthetic Gene Cluster
ESKAPE: *Enterococcus faecium, Staphylococcus aureus, Klebsiella pneumoniae, Acinetobacter baumannii, Pseudomonas aeruginosa,* and *Enterobacter spp*.
FBMN: Feature-Based Molecular Networking
GNPS: Global Natural Products Social Molecular Networking
LC-MS/MS: Liquid Chromatography-Tandem Mass Spectrometry
MRSA: Methicillin-Resistant *Staphylococcus aureus*
NP: Natural Product
NRPS: Nonribosomal Peptide Synthetase
OSMAC: One Strain Many Compounds PKS Polyketide Synthase
QC: Quality Control
SCA: Starch-Casein Agar
SCB: Starch-Casein Broth
UHPLC-HRMS/MS: Ultra-High-Performance Liquid Chromatography-High-Resolution Tandem Mass Spectrometry

## Notes

https://doi.org/10.25345/C5XP6VJ31

https://gnps2.org/status?task=a2721f67b9af435da7be0db18b25fbae

https://www.ebi.ac.uk/ena/browser/view/PRJEB108104

### Competing Interest Statement

The authors have declared no competing interest.

