## Supplementary Figures and Tables for "An Integrated Miniaturized Metabologenomics Workflow for High-Throughput Actinomycetes Prioritization": Supplementary_Figures.pdf

**Supplementary Figure S1. GNPS mirror-spectrum validation of Actinomycin D and Actinomycin X2.** Mirror-spectrum comparisons supporting the annotation of Actinomycin D (features 3330 and 3333) and Actinomycin X2 (features 3327 and 3329). High spectral similarity between experimental and library spectra further supports the assignment of the major congeners comprising the actinomycin molecular family.

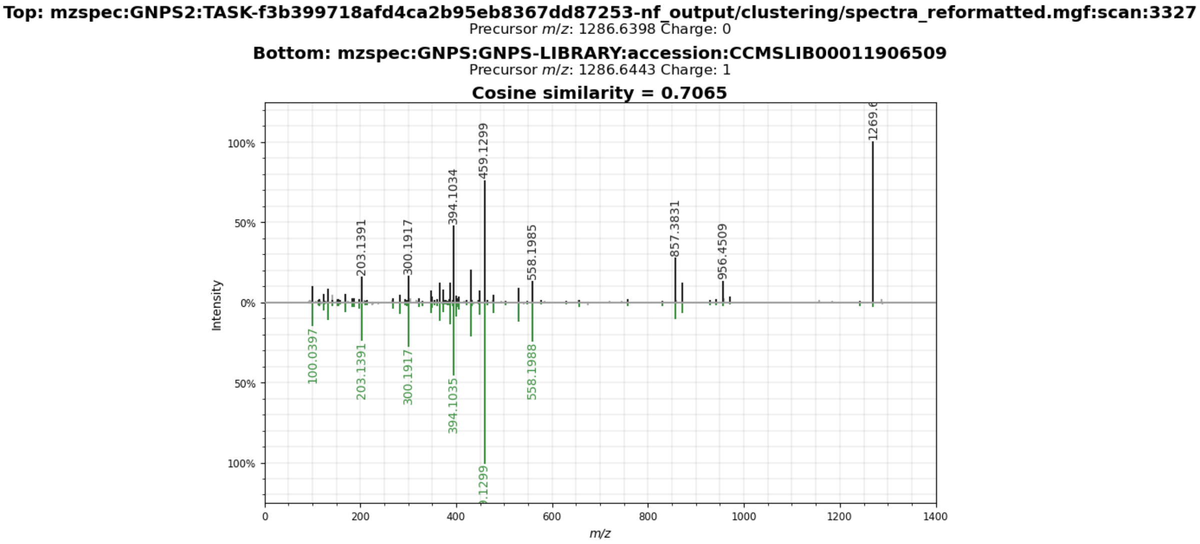


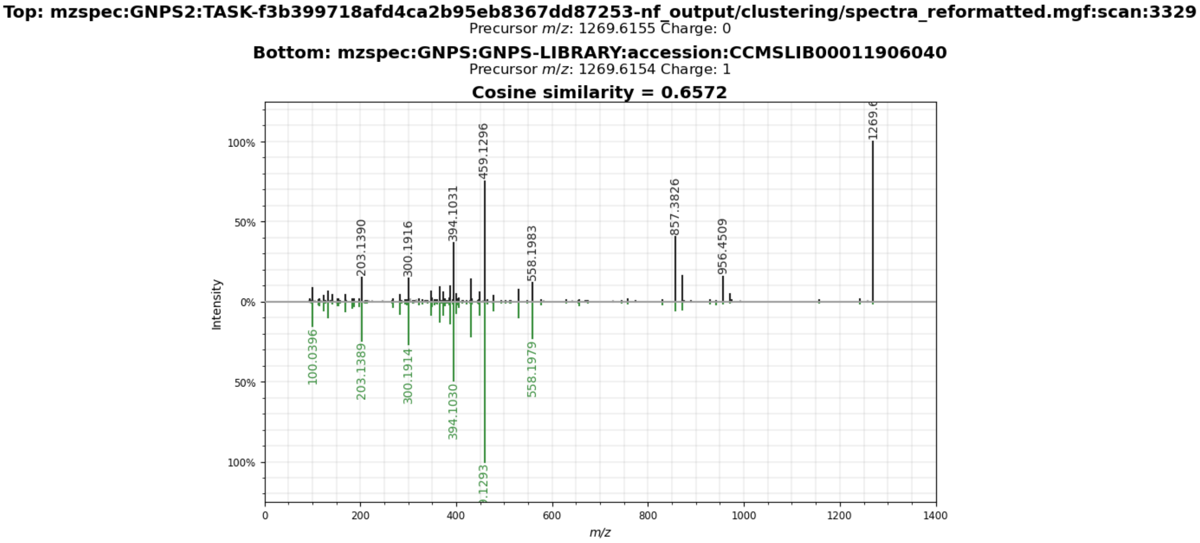

**Top: mzspec:GNPS2:TASK-f3b399718afd4ca2b95eb8367dd87253-nf_output/clustering/spectra_reformatted.mgf:scan:3330**
Precursor *m/z*: 1255.6380 Charge: 0

**Bottom: mzspec:GNPS:GNPS-LIBRARY:accession:CCMSLIB00011906039**
Precursor *m/z*: 1255.6367 Charge: 1

**Cosine similarity = 0.8084**

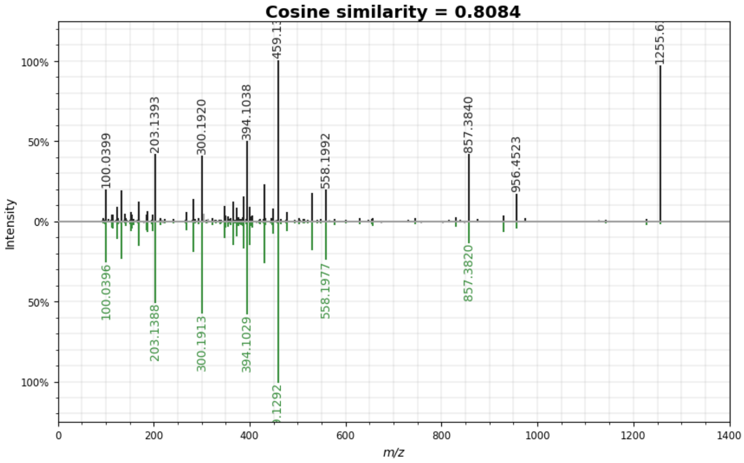


**Top: mzspec:GNPS2:TASK-f3b399718afd4ca2b95eb8367dd87253-nf_output/clustering/spectra_reformatted.mgf:scan:3333**
Precursor *m/z*: 1277.6177 Charge: 0

**Bottom: mzspec:GNPS:GNPS-LIBRARY:accession:CCMSLIB00011906328**
Precursor *m/z*: 1277.6200 Charge: 1

**Cosine similarity = 0.8145**

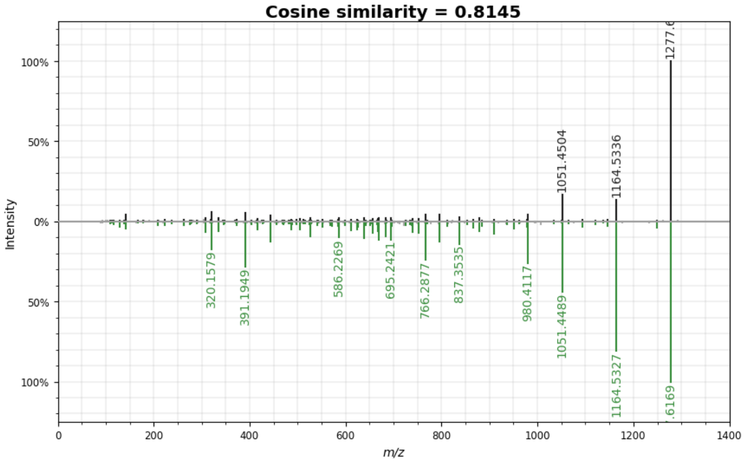

**Supplementary Figure S2. GNPS mirror-spectrum validation of Nigericin.** Mirror-spectrum comparison of feature 3478 (*m/z* 747.4637), annotated as Nigericin, showing a cosine similarity score of 0.987 against the GNPS reference spectrum and providing high-confidence support for its annotation within the polyether ionophore molecular family.

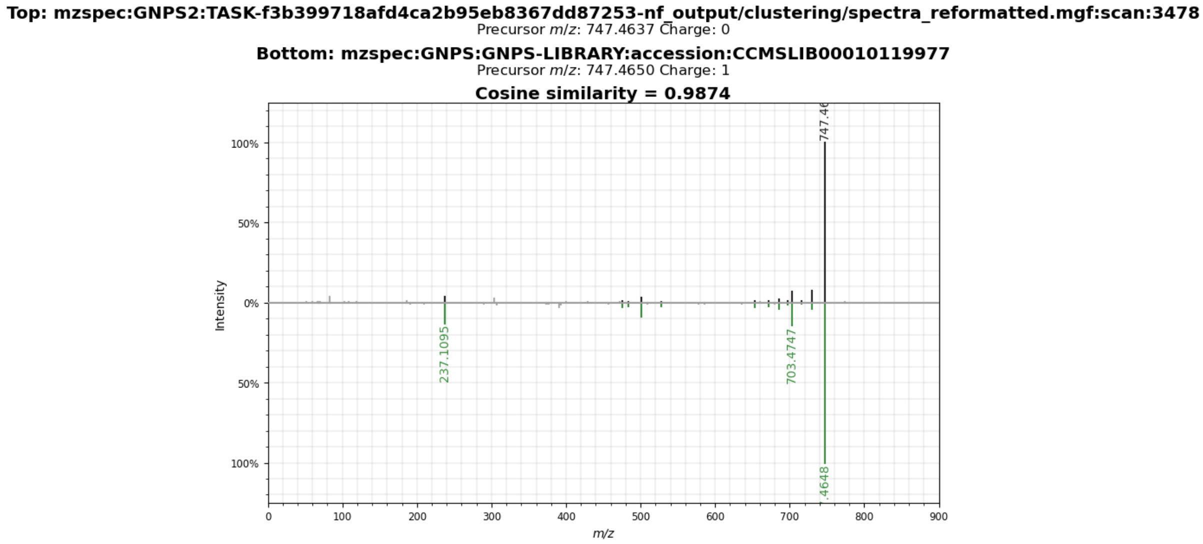


**Supplementary Figure S3. GNPS mirror-spectrum validation of Fridamycin B.** Mirror-spectrum comparison of feature 3016 (*m/z* 487.1602) against the GNPS reference spectrum of Fridamycin B [M+H]+. The spectral similarity score (0.725) supports the assignment of Component 108 to a fridamycin-related angucycline molecular family.

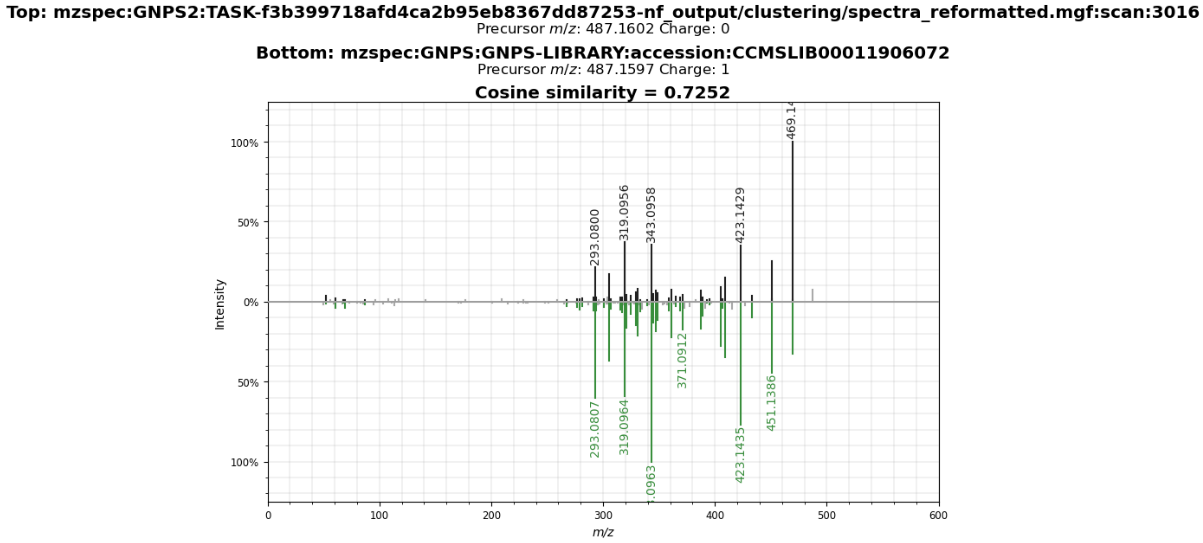

**Supplementary Figure S4. GNPS mirror-spectrum validation of Geldanamycin.** Mirror-spectrum comparison of feature 3230 (*m/z* 583.2643) against the GNPS reference spectrum of Geldanamycin [M+Na]+. Although the spectral similarity score was moderate, complementary analyses from GNPS, ISDB-Lotus, SIRIUS, CANOPUS, FBMN, and genome mining support the annotation, all consistent with the production of ansamycin metabolites.

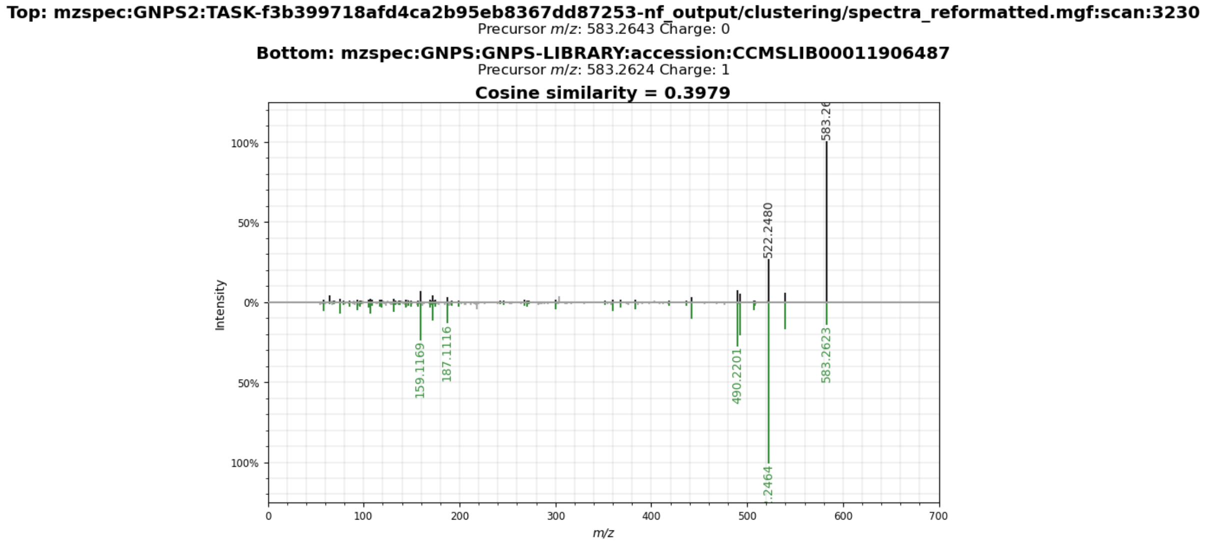

**Supplementary Figure S5. GNPS mirror-spectrum validation of ectoine-derived compatible solutes.** Mirror-spectrum comparison of feature 214 (m/z 143.0816 [M+H]+) and feature 15 (m/z 159.0766 [M+H]+) against the GNPS reference spectra of ectoine and 5-hydroxyectoine, respectively. The high spectral similarity observed for both metabolites (cosine scores = 0.899 and 0.9995, respectively), together with complementary evidence from ISDB-Lotus, SIRIUS, CANOPUS, and FBMN, supports the assignment of Component 145 to an ectoine-associated molecular family.

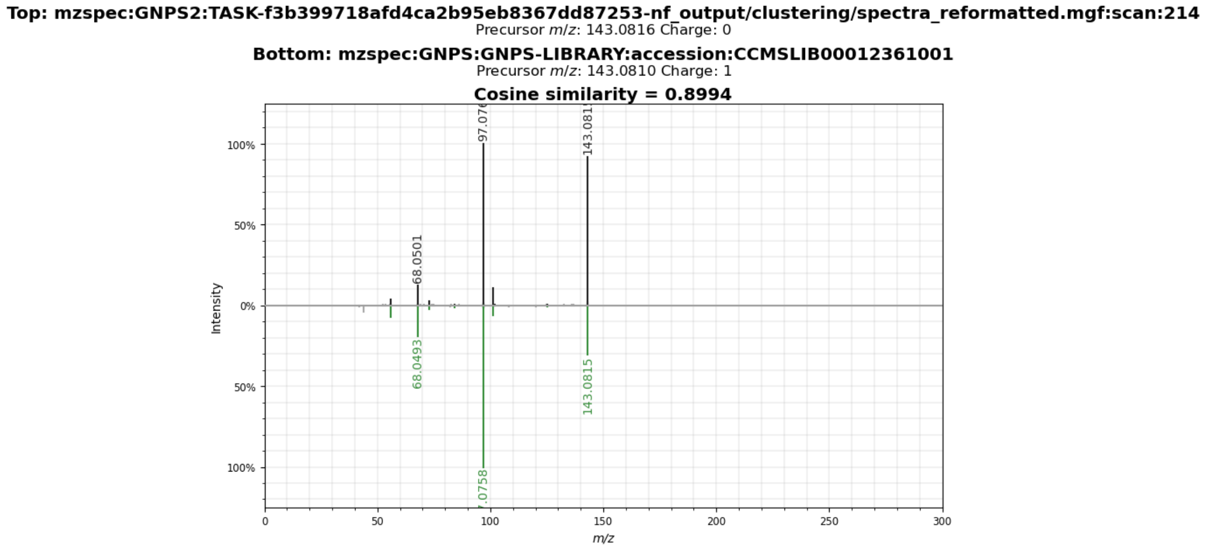


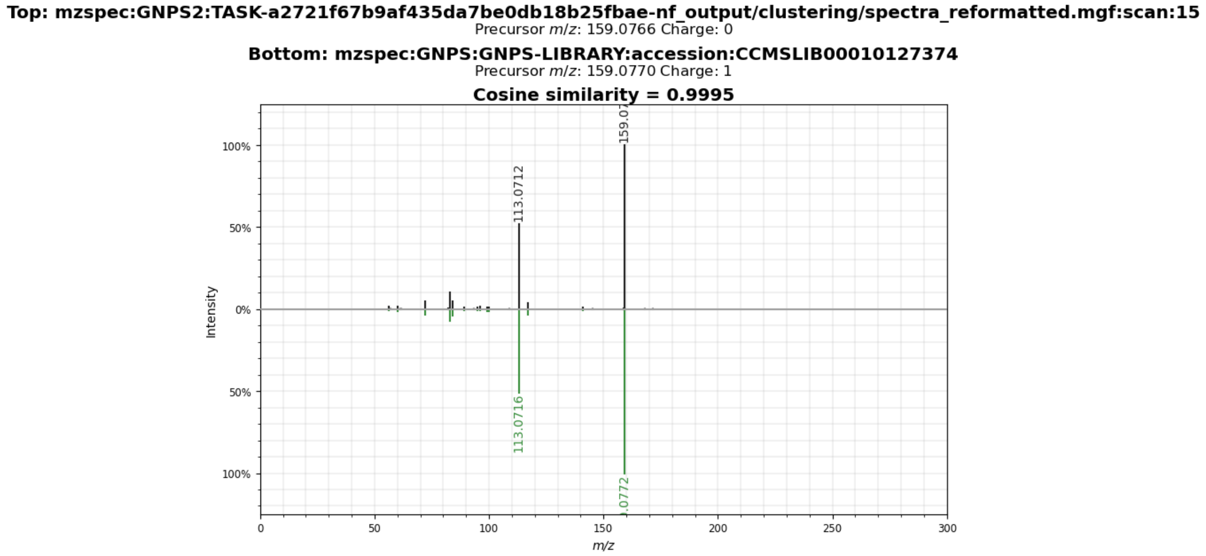

**Supplementary Figure S6. GNPS mirror-spectrum validation of lydicamycin-related metabolites.** Mirror-spectrum comparisons of features 2501 (*m/z* 841.5328), annotated as TPU-0037A (lydicamycin derivative), and 2583 (*m/z* 853.5322), annotated as a putative lydicamycin analogue. The high spectral similarity scores (0.891 and 0.930, respectively) provide additional support for the assignment of component 79 to a lydicamycin-related molecular family.

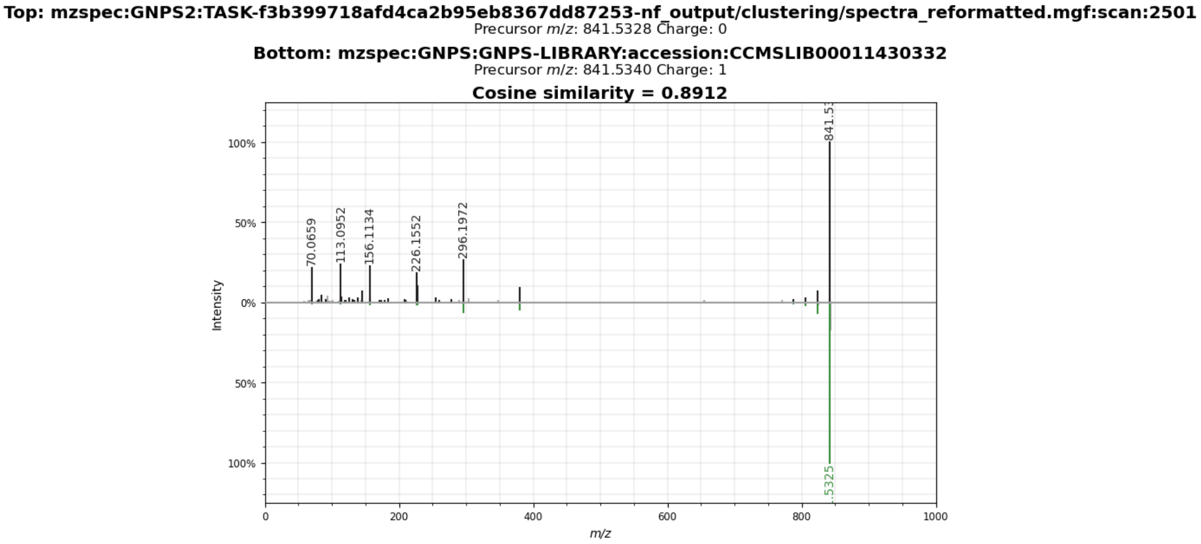


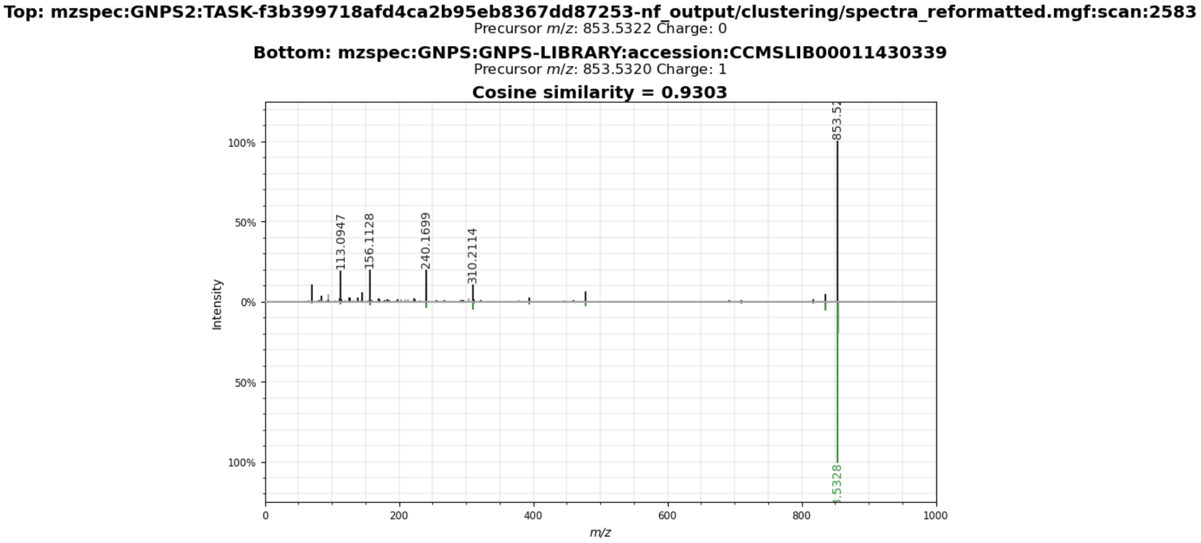

**Supplementary Figure S7. GNPS mirror-spectrum validation of Tambromycin A.** Mirror-spectrum comparison of feature 2473 (*m/z* 536.1915 [M+H]+) against the GNPS reference spectrum of Tambromycin A (CCMSLIB00006112534). The observed spectral similarity (cosine score = 0.7021) supports annotating Component 44 as a tambromycin-associated molecular family.

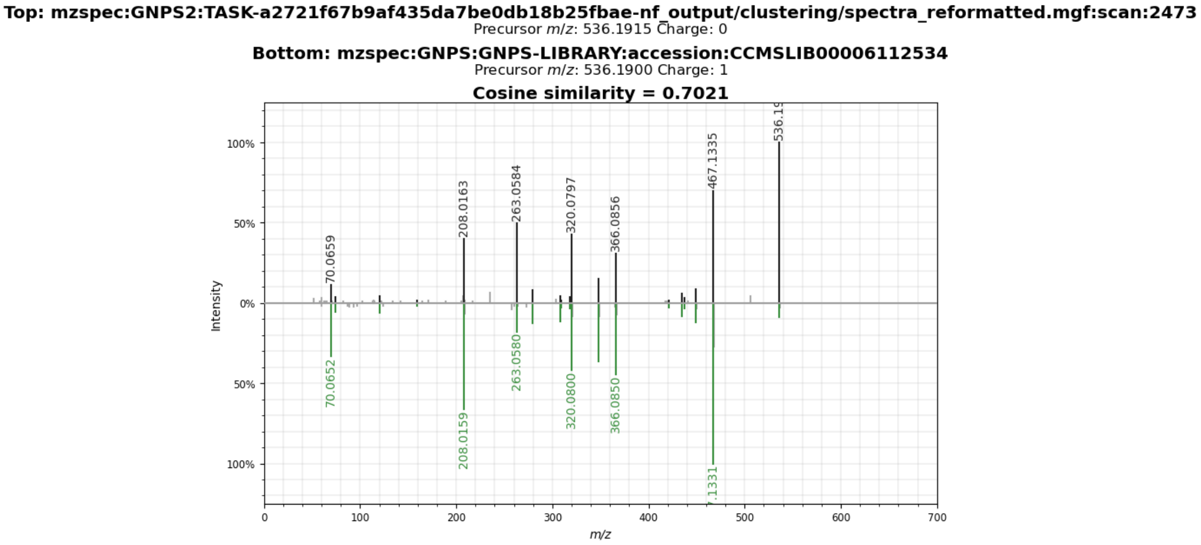

**Supplementary Figure S8.** Representative chromatographic comparisons used to support selected metabolite annotations. The figure shows extracted ion chromatogram (EIC) overlays of authentic standards, corresponding LC-MS features detected in pooled QC samples or individual culture extracts, and extraction blank controls for actinomycin D, ectoine, 5-hydroxyectoine, geldanamycin, monensin A, and nigericin. Detailed comparisons are provided in Supplementary Table S6.

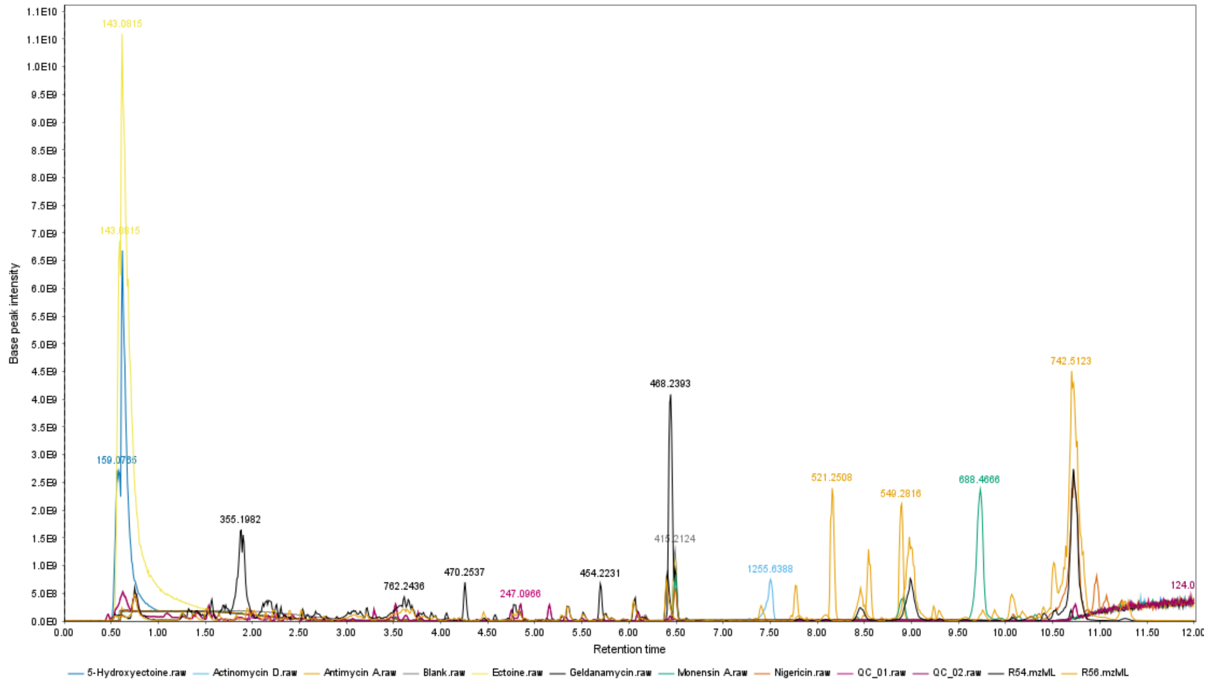

### S8A. Actinomycin D

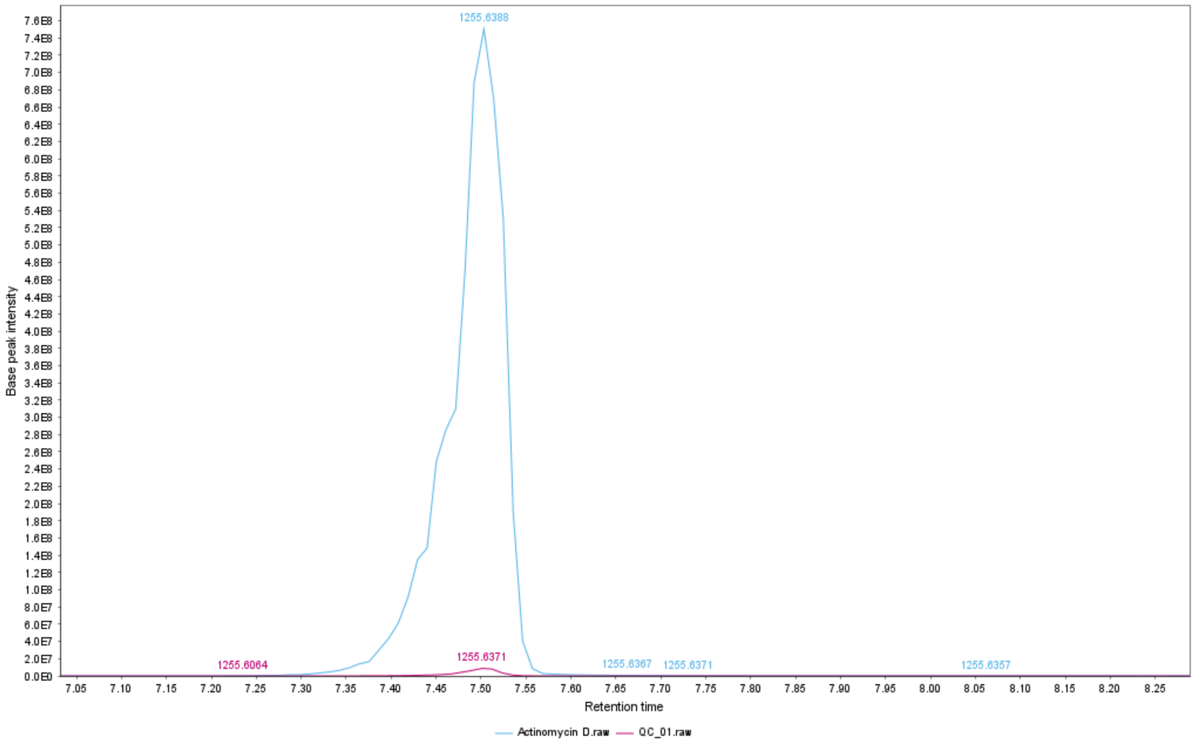


### S8B. Nigericin [M+Na]

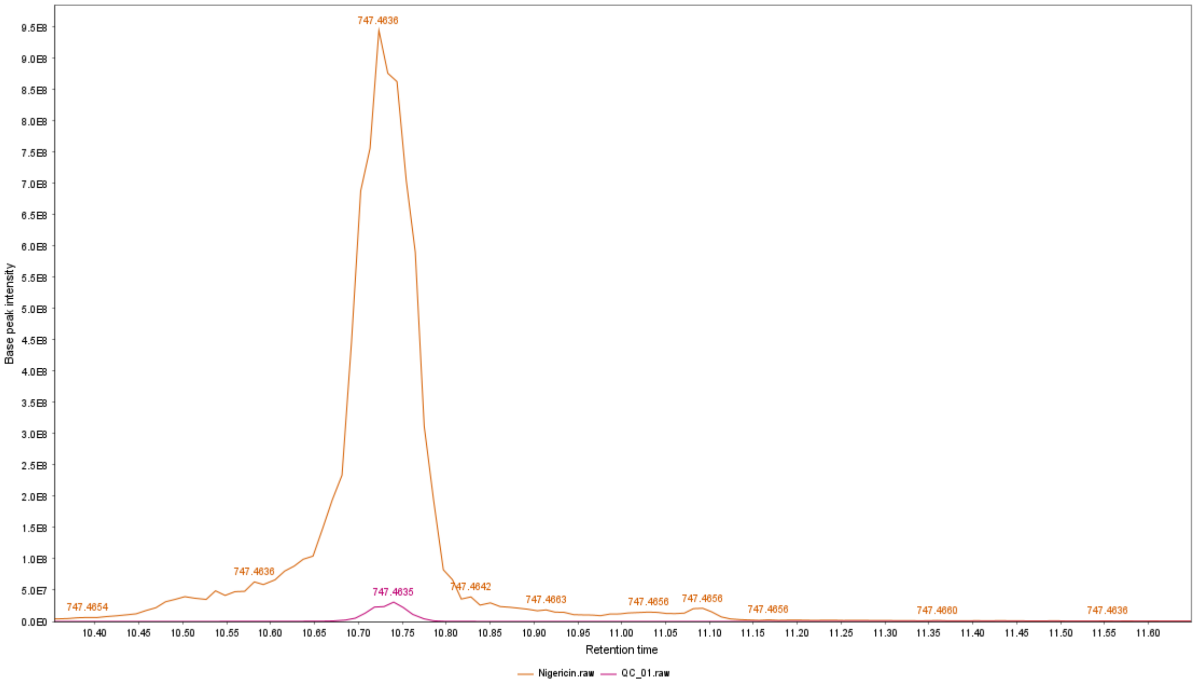

### S8C. Monensin A

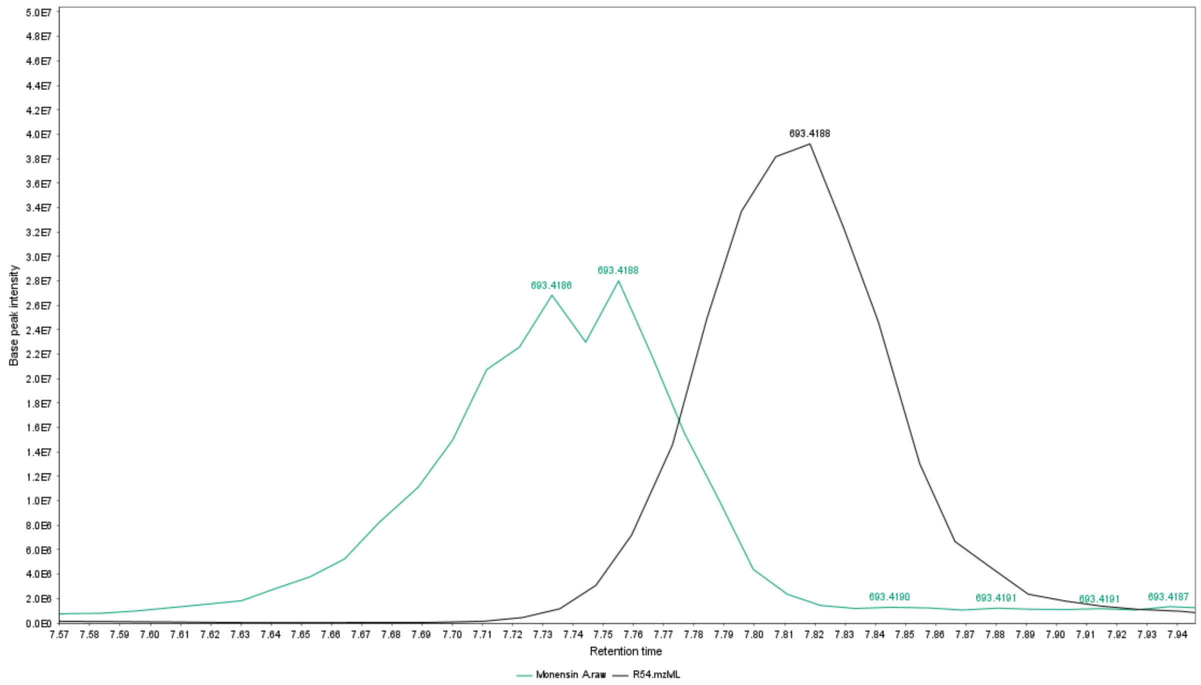


### S8D. Geldanamycin

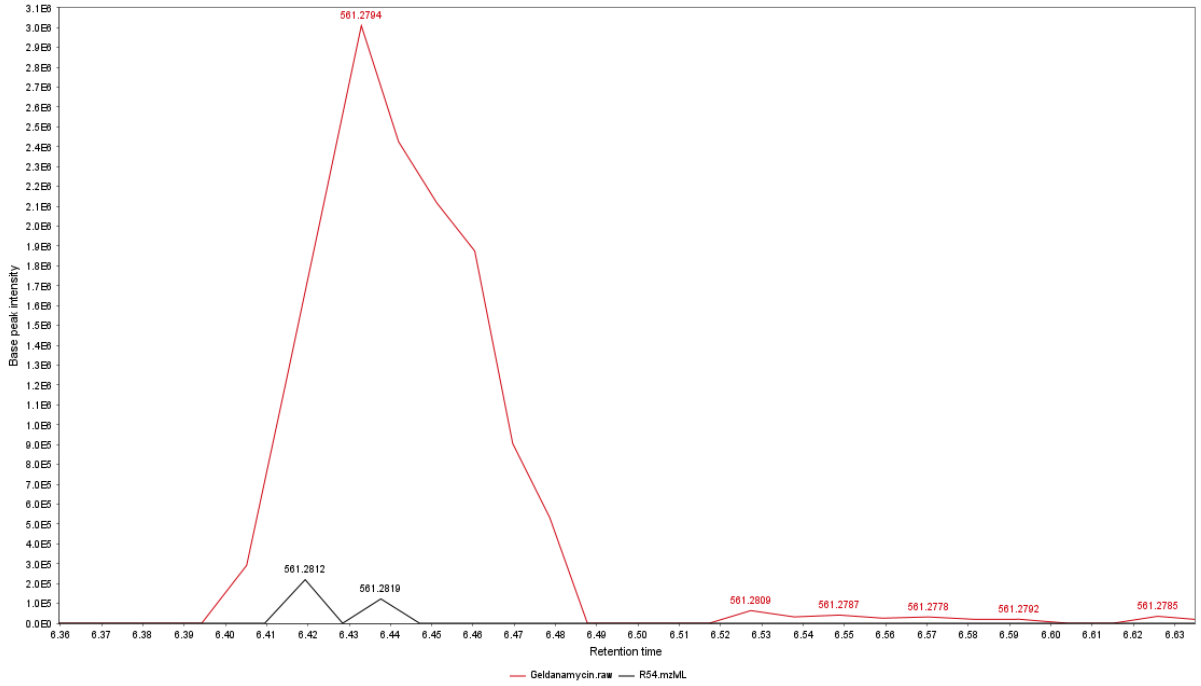

### S8E. Ectoine

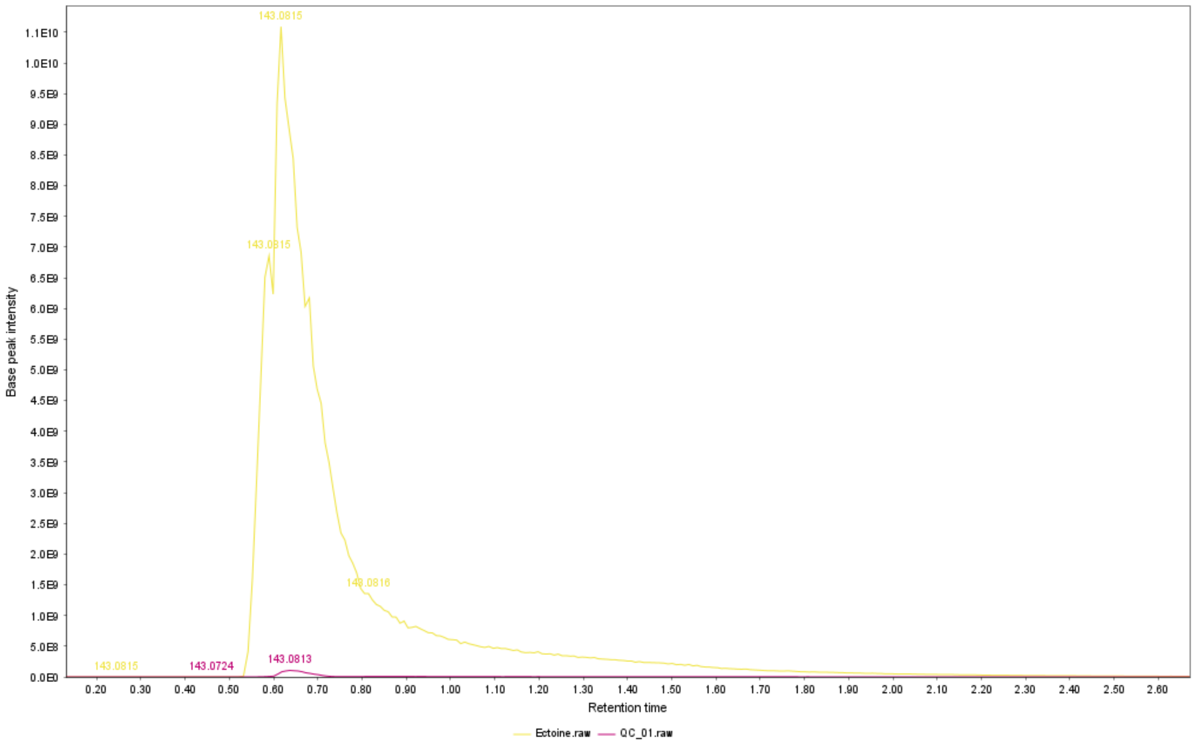


### S8F. 5-Hydroxyectoine

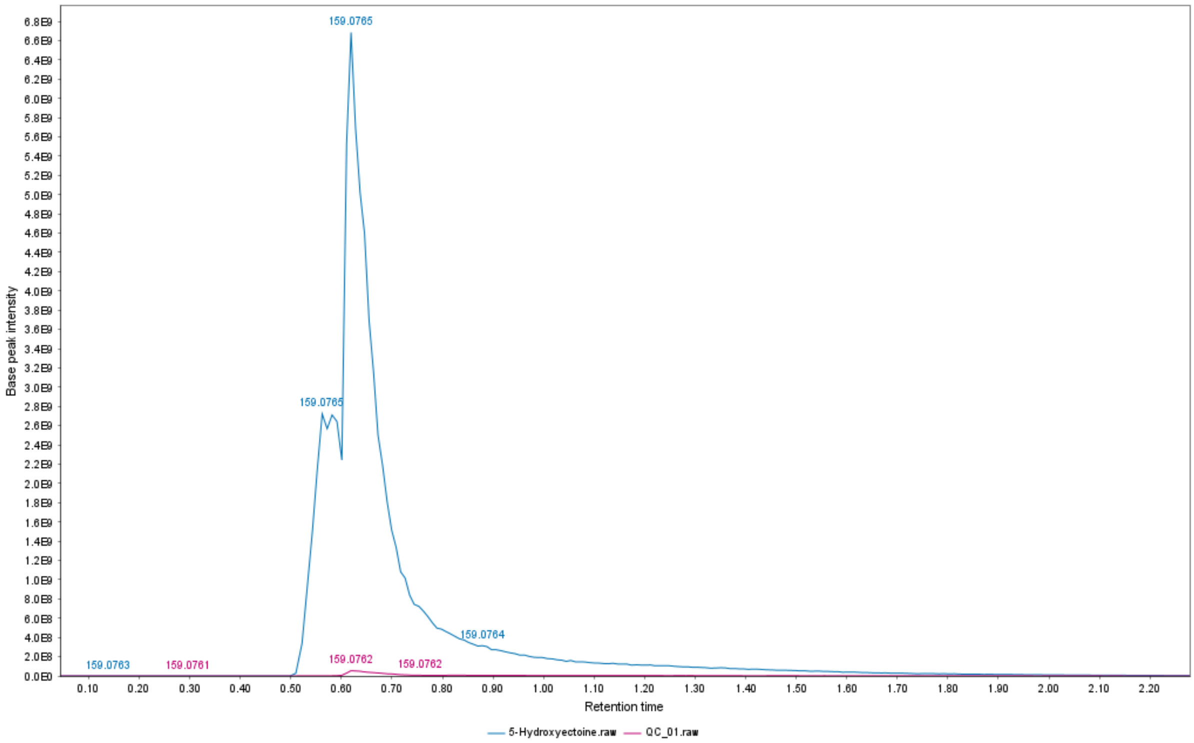
